# Regulation of positive and negative selection and TCR signaling during thymic T cell development by capicua

**DOI:** 10.1101/2021.07.11.451936

**Authors:** Soeun Kim, Guk-Yeol Park, Jong Seok Park, Jiho Park, Hyebeen Hong, Yoontae Lee

**Author notes:** Correspondence to: Yoontae Lee, Room 388, POSTECH Biotech Center, 77 Cheongam-Ro, Nam-Gu, Pohang, Gyeongbuk 37673, Republic of Korea.

## Abstract

Central tolerance is achieved through positive and negative selection of thymocytes mediated by T cell receptor (TCR) signaling strength. Thus, the dysregulation of the thymic selection process often leads to autoimmunity. Here, we show that capicua (CIC), a transcriptional repressor that suppresses autoimmunity, controls the thymic selection process. Loss of CIC prior to T-cell lineage commitment impaired both positive and negative selection of thymocytes. CIC deficiency attenuated TCR signaling in CD4^+^CD8^+^ double-positive (DP) cells, as evidenced by a decrease in CD5 and phospho-ERK levels and calcium flux. We identified *Spry4*, *Dusp4*, *Dusp6,* and *Spred1* as CIC target genes that could inhibit TCR signaling in DP cells. Furthermore, impaired positive selection and TCR signaling were partially rescued in *Cic* and *Spry4* double mutant mice. Our findings indicate that CIC is a transcription factor required for thymic T cell development and suggest that CIC acts at multiple stages of T cell development and differentiation to prevent autoimmunity.

## INTRODUCTION

T cells play a crucial role in the adaptive immune system’s defense against external invasion. To distinguish between self and non-self, T cells use their T cell receptors (TCRs) to recognize peptide-loaded major histocompatibility complex (MHC) molecules and respond to many types of antigens with an enormous TCR repertoire. T cells with a specific TCR are generated in the thymus through a series of processes, ranging from random rearrangement of TCR gene segments to selection processes that entail apoptosis of inappropriate cells (Klein, Kyewski, Allen, & Hogquist, 2014). T cell development in the thymus is tightly regulated to prevent the generation of nonfunctional or self-reactive T cells. Progenitor cells from bone marrow (BM) become CD4^-^CD8^-^ double-negative (DN) cells and undergo a process called β-selection, which selects only T cells with a functional TCRβ chain. DN cells that have passed β-selection then become CD4^+^CD8^+^ double-positive (DP) cells and undergo positive selection, which selects T cells that bind to self-peptide ligands loaded onto MHC (self-pMHC) molecules. These CD4^+^CD8^+^ DP cells then develop into CD4^+^CD8^-^ single-positive (CD4^+^ SP) or CD4^-^CD8^+^ single-positive (CD8^+^ SP) T cells. Negative selection, also known as clonal deletion, selectively removes T cells that bind with high affinity to self-pMHC during the DP and SP stages. Because the intensity and duration of TCR signaling based on TCR affinity to self-pMHC are the major determinants of selection (Gascoigne, Rybakin, Acuto, & Brzostek, 2016), defects in the TCR signaling component lead to abnormal T cell development and alteration of the TCR repertoire (Guoping Fu et al., 2010; Sakaguchi et al., 2003). Impairment of thymic selection caused by decreased TCR signaling destroys central tolerance, and consequently induces autoimmunity accompanied by the expansion of the CD44^hi^CD62L^lo^ activated effector/memory T cell population (Guoping Fu et al., 2010; Sakaguchi et al., 2003; Sommers et al., 2002).

Capicua (CIC) is an evolutionarily conserved transcriptional repressor that regulates the receptor tyrosine kinase (RTK) signaling pathway in *Drosophila* and mammals (Jiménez, Guichet, Ephrussi, & Casanova, 2000; Jiménez, Shvartsman, & Paroush, 2012). CIC is expressed as two different isoforms: long (CIC-L) and short (CIC-S), which differ in their amino termini (Y. Lee, 2020). CIC recognizes specific octameric DNA sequences (5ʹ-T(G/C)AATG(A/G)(A/G)-3ʹ) within its target gene promoter regions and represses their expression (Kawamura-Saito et al., 2006; Shin & Hong, 2014; Weissmann et al., 2018). Several genomic and transcriptomic analyses have identified CIC target genes, including *ETV1*, *ETV4*, *ETV5*, *SPRY4*, *SPRED1*, *DUSP4*, and *DUSP6*, in various types of cells (Fryer et al., 2011; Weissmann et al., 2018; Yang et al., 2017). Activation of the RTK/RAS/MAPK signaling pathway phosphorylates and inactivates CIC via dissociation from target gene promoters, cytoplasmic translocation, and/or proteasomal degradation (Jiménez et al., 2012; Keenan et al., 2020; Y. Lee, 2020). CIC also mediates the ERK-DUSP6 negative feedback loop to maintain ERK activity within the physiological range in mammals (Ren et al., 2020).

We previously reported that murine CIC deficiency spontaneously induces lymphoproliferative autoimmune-like phenotypes (S. Park et al., 2017). Hematopoietic lineage cell-specific (*Vav1-Cre*-mediated knockout) and T cell-specific (*Cd4-Cre*-mediated knockout) *Cic*-null mice commonly exhibit autoimmune-like symptoms including hyperglobulinemia, increased serum anti-dsDNA antibody levels, and tissue infiltration of immune cells, accompanied by increased frequency of CD44^hi^CD62L^lo^ T and follicular helper T (Tfh) cells in the spleen (S. Park et al., 2017). However, these phenotypes were more severe in *Cic^f/f^;Vav1-Cre* mice than in *Cic^f/f^;Cd4-Cre* mice; moreover, enlargement of secondary lymphoid organs was observed in *Cic^f/f^;Vav1-Cre* mice but not in *Cic^f/f^;Cd4-Cre* mice (G. Park et al., 2020; S. Park et al., 2017). These results indicated that deletion of *Cic* alleles in hematopoietic stem and progenitor cells leads to more severe peripheral T cell hyperactivation and autoimmunity than the *Cd4-Cre*-mediated *Cic* deletion in CD4^+^CD8^+^ DP thymocytes. We also reported that enhanced peripheral T cell hyperactivation in *Cic^f/f^;Vav1-Cre* mice relative to *Cic^f/f^;Cd4-Cre* mice was caused by CIC deficiency in T cells rather than in other types of immune cells (G. Park et al., 2020), suggesting that this abnormality could be caused by impaired control of early thymic T cell development in *Cic^f/f^;Vav1-Cre* mice. It was reported that the frequency of CD4^-^CD8^-^CD44^+^CD25^-^ DN1 cells was increased in adult stage-specific *Cic-*null mice (Tan et al., 2018). Taken together, these studies suggest that CIC plays a crucial role in thymic T cell development.

In this study, we found that CIC regulates thymic T cell development from the CD4^-^ CD8^-^ DN stage and positive and negative selection of thymocytes, primarily during the CD4^+^CD8^+^ DP stage. TCR signaling was significantly attenuated in DP cells of *Cic^f/f^;Vav1-Cre* mice, thereby impairing both positive and negative selection in *Cic^f/f^;Vav1-Cre* mice. We also identified *Spry4*, *Dusp4*, *Dusp6,* and *Spred1* as CIC target genes that could potentially contribute to the reduced TCR signaling strength and impaired thymic selection processes in *Cic^f/f^;Vav1-Cre* mice. Our findings demonstrate that CIC is a critical regulator of TCR signaling in DP cells and thymic T cell development.

## RESULTS

### Changes in the frequencies of thymic T cell subsets over development in *Cic^f/f^;Vav1-Cre* mice

To examine the role of CIC in thymic T cell development, we first evaluated the levels of CIC in multiple subsets of developing thymocytes using homozygous FLAG-tagged *Cic* knock-in (*Cic^FLAG/FLAG^*) mice (S. Park, Park, Kim, & Lee, 2019). Flow cytometry for FLAG-CIC revealed that CIC levels were relatively high in CD4^-^CD8^-^ DN and immature CD8^+^ single positive (ISP) cells compared to cells at later developmental stages, such as CD4^+^CD8^+^ DP and CD4^+^ or CD8^+^ SP subsets (Fig. 1A). We then determined the frequency and number of thymic T cell subsets in *Cic^f/f^* (WT), *Cic^f/f^;Cd4-Cre,* and *Cic^f/f^;Vav1-Cre* mice at 7 weeks of age. The total numbers of thymocytes were comparable among WT, *Cic^f/f^;Cd4-Cre,* and *Cic^f/f^;Vav1-Cre* mice (Fig. 1B). However, T cell development during the DN stage was abnormal in *Cic^f/f^;Vav1-Cre* mice, as evidenced by an increased frequency of CD44^hi^CD25^hi^ DN2 and CD44^lo^CD25^hi^ DN3 subsets at the expense of the CD44^lo^CD25^lo^ DN4 subset (Fig. 1C). As expected, these changes were not detected in *Cic^f/f^;Cd4-Cre* mice (Fig. 1C), because *Cd4-Cre* was not expressed in DN cells (P. P. Lee et al., 2001). We previously reported that the frequency of DN, DP, and SP cells was comparable between WT and *Cic^f/f^;Vav1-Cre* mice at 9 weeks of age (S. Park et al., 2017). However, at 7 weeks of age, *Cic^f/f^;Vav1-Cre* mice exhibited a mild block in formation of SP cells in the thymus: the frequency of DP cells was significantly increased in *Cic^f/f^;Vav1-Cre* mice compared to WT and *Cic^f/f^;Cd4-Cre* mice, whereas that of SP cells decreased (Fig. 1D). This difference was not statistically significant when calculated using cell numbers (Fig. 1D). To clarify whether CIC deficiency affected thymic SP cell formation, we performed the same analysis using WT and *Cic^f/f^;Vav1-Cre* mice at 1 week of age, when the proportion of peripheral T cells that recirculated into the thymus was negligible (Hale, Boursalian, Turk, & Fink, 2006). The frequency and number of CD4^+^ and CD8^+^ SP cells were significantly decreased in 1-week-old *Cic^f/f^;Vav1-Cre* mice (Fig. 1E), suggesting that CIC regulates SP cell development in the thymus. Consistent with this result, both CD4^+^ and CD8^+^ T cell populations were substantially reduced in the spleens of 1-week-old *Cic^f/f^;Vav1-Cre* mice (Fig. S1). Together, these data demonstrate that CIC is involved in the regulation of thymic T cell development.

**Figure 1.**
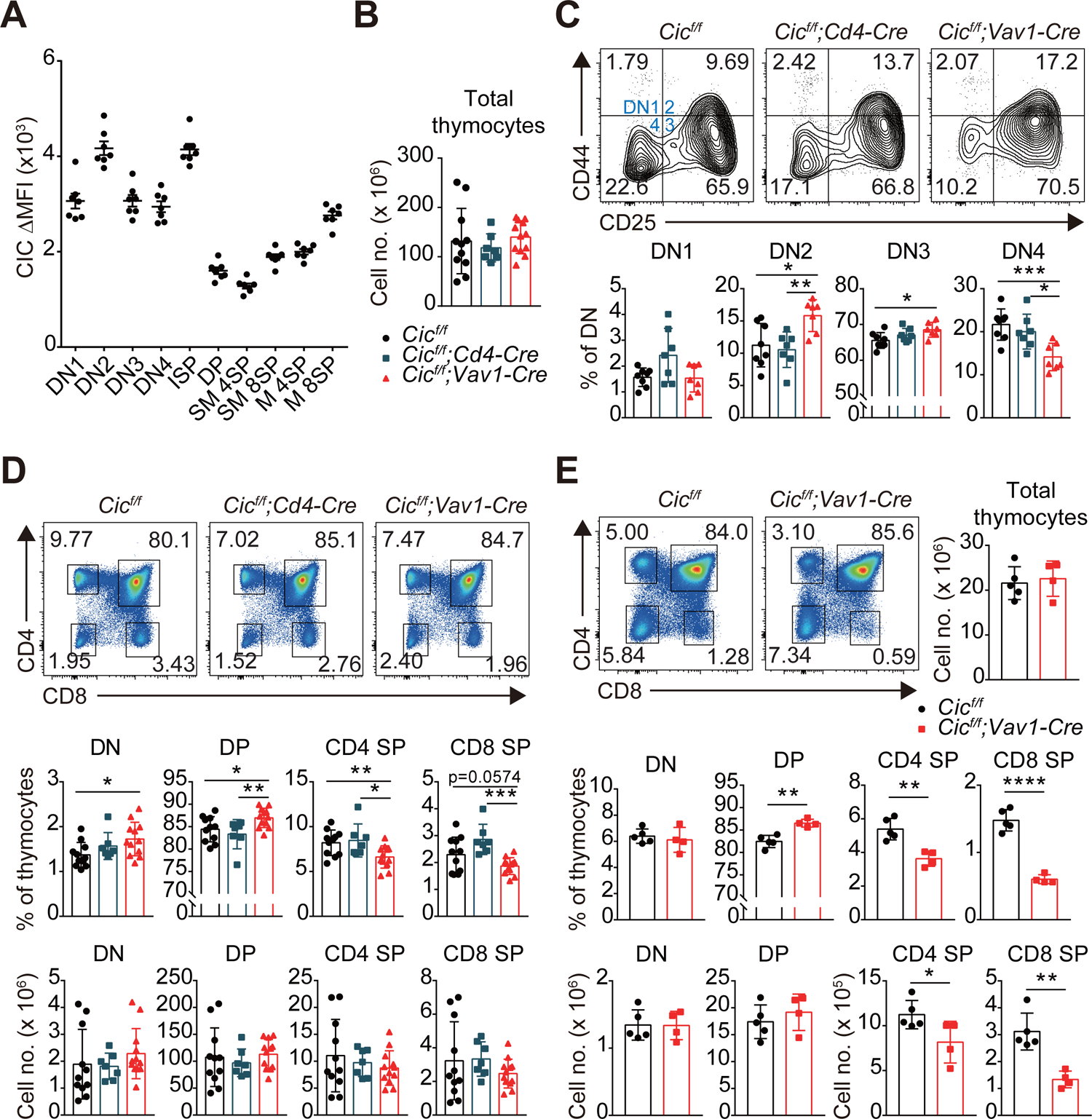
Altered T cell development in *Cic^f/f^;Vav1-cre* mice. **(A)** CIC protein levels in thymic T cell subsets. Thymocytes of *Cic^FLAG/FLAG^* mice (N = 7) were subjected to flow cytometry using anti-FLAG antibody. The ΔMFI of CIC-FLAG was calculated by subtraction of the mean fluorescence intensity (MFI) value obtained by isotype control from that obtained by the anti-FLAG antibody. DN1: CD4^-^CD8^-^CD44^hi^CD25^lo^, DN2: CD4^-^CD8^-^CD44^hi^CD25^hi^, DN3: CD4^-^CD8^-^CD44^lo^CD25^hi^, DN4: CD4^-^CD8^-^CD44^lo^CD25^lo^; ISP: immature CD8^+^ single positive cells (CD4^-^CD8^-^TCRb^lo^CD24^hi^), DP: CD4^+^CD8^+^, SM: semi-mature (CD69^+^TCRb^hi^), and M: mature (CD69^-^TCRb^hi^). **(B-D)** Flow cytometry of thymocytes from 7-week-old *Cic^f/f^*, *Cic^f/f^;Cd4-Cre*, and *Cic^f/f^;Vav1-Cre* mice. **(B)** Total numbers of thymocytes for each genotype. N = 11, 7, and 12 for *Cic^f/f^*, *Cic^f/f^;Cd4-Cre*, and *Cic^f/f^;Vav1-Cre* mice, respectively. **(C)** Proportions of DN1-4 subsets for each genotype. N = 8, 7, and 7 for *Cic^f/f^*, *Cic^f/f^;Cd4-Cre*, and *Cic^f/f^;Vav1-Cre* mice, respectively. **(D)** Frequencies and numbers of DN, DP, CD4^+^ SP, and CD8^+^ SP cells for each genotype. N = 11, 7, and 12 for *Cic^f/f^*, *Cic^f/f^;Cd4-Cre*, and *Cic^f/f^;Vav1-Cre* mice, respectively. **(E)** Flow cytometry of thymocytes of 1-week-old *Cic^f/f^* and *Cic^f/f^;Vav1-Cre* mice using CD4 and CD8 markers. Total thymocyte numbers, and the frequencies and numbers of DN, DP, CD4 SP, and CD8 SP subsets in mice of each genotype, as well as representative plot images, are presented. N = 5 and 4 for *Cic^f/f^* and *Cic^f/f^;Vav1-Cre* mice, respectively. Data are representative of two independent experiments. Bar graphs show data as means with SEM. *P < 0.05, **P < 0.01, ***P < 0.001, and ****P < 0.0001. The unpaired two-tailed Student’s t-test was used to calculate P values. See also Figure source data 1.

### Stable CIC expression in DP cells of *Cic^f/f^;Cd4-Cre* mice

After obtaining the results shown in Fig. 1D, we wondered why the frequency of DP cells was comparable between WT and *Cic^f/f^;Cd4-Cre* mice because *Cic* alleles were supposed to be deleted in DP cells by *Cd4-Cre* (P. P. Lee et al., 2001). Western blotting for CIC in multiple developing thymic T cell subsets from WT, *Cic^f/f^;Cd4-Cre,* and *Cic^f/f^;Vav1-Cre* mice provided the answer. Although CIC expression disappeared in all tested thymic T cell subsets of *Cic^f/f^;Vav1-Cre* mice, CIC was still substantially expressed in the DP cells of *Cic^f/f^;Cd4-Cre* mice (Fig. 2A and Fig. S4C). This appeared to be caused by CIC protein stability, because the *loxP* site-flanked genomic regions containing exons 9–11 of *Cic* were completely removed in DP cells by *Cd4-Cre* and *Vav1-Cre* (Fig. 2B). These data suggest that the impaired SP cell formation in the thymus of *Cic^f/f^;Vav1-Cre* mice is attributed to the loss of CIC in DN and/or DP cells rather than that in SP cells, because CIC expression was not detected in SP thymocytes from *Cic^f/f^;Cd4-Cre* and *Cic^f/f^;Vav1-Cre* mice (Fig. 2A and Fig. S4C).

**Figure 2.**
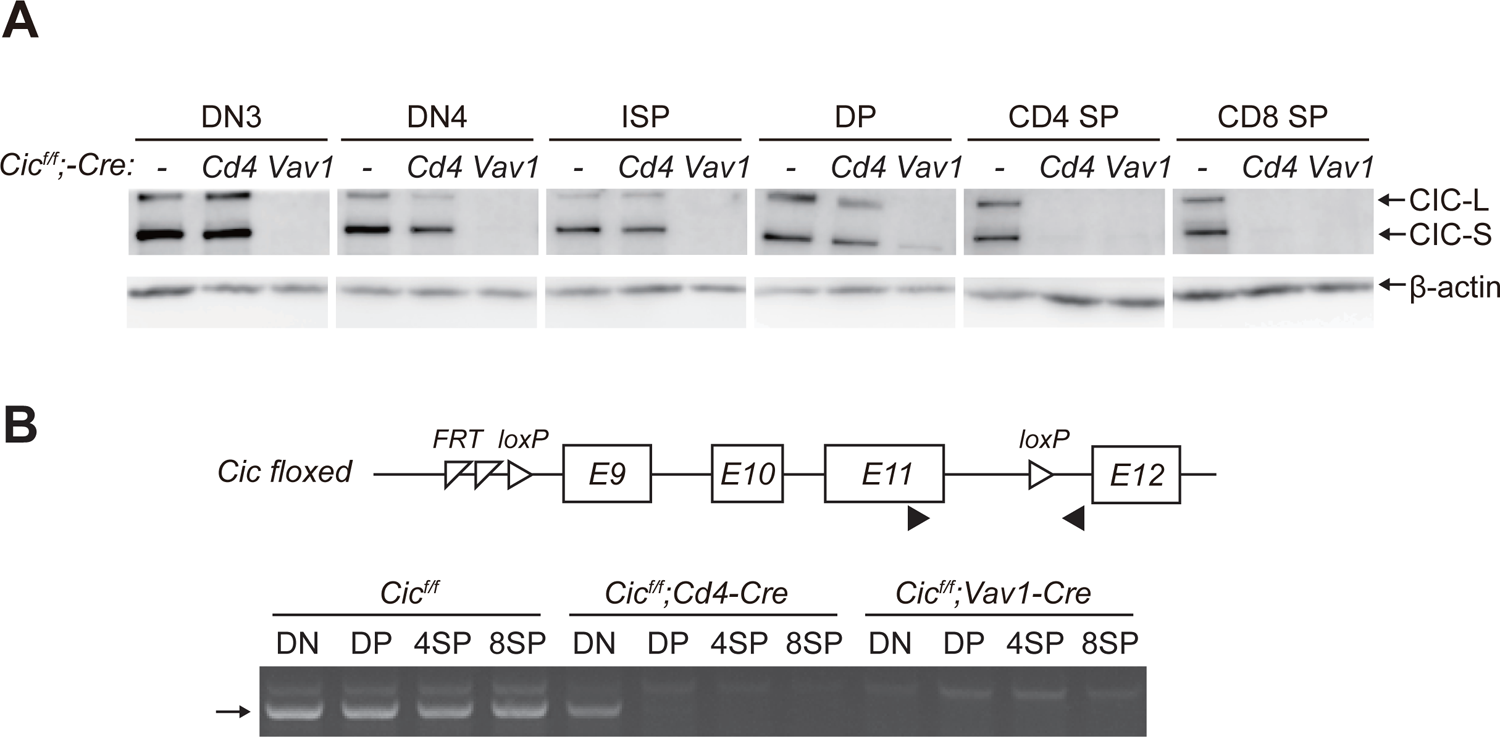
Stable CIC expression in DP cells of *Cic^f/f^;Cd4-Cre* mice. **(A)** Western blotting for CIC levels in DN3, DN4, ISP, DP, CD4^+^ SP, and CD8^+^ SP cells from *Cic^f/f^*, *Cic^f/f^;Cd4-Cre*, and *Cic^f/f^;Vav1-Cre* mice. Lin^-^ (Lineage (CD11b, CD11c, CD19, NK1.1, Gr-1, γδTCR, and TER119)-negative) gated DN3, DN4, ISP, DP, CD4^+^ SP, and CD8^+^ SP cells were sorted from mice of each genotype. 1-PCR analysis of Cic knock-out efficiency in DN, DP, CD4^+^ SP, and CD8^+^ SP cells from *Cic^f/f^*, *Cic^f/f^;Cd4-Cre*, and *Cic^f/f^;Vav1-Cre* mice. Genomic DNA was extracted from sorted DN, DP, CD4^+^ SP, and CD8^+^ SP cells and subjected to PCR for a part of Cic floxed allele. Upper panel, schematic of the Cic floxed allele. Arrowheads indicate the primers used for PCR. Lower panel, representative agarose gel image of PCR bands. An arrow indicates PCR bands of the Cic floxed allele. See also Figure source data 1.

### Impaired thymic positive selection in *Cic^f/f^;Vav1-Cre* mice

Defects in thymic positive selection that occurs during the CD4^+^CD8^+^ DP developmental stage often result in a decrease in the CD4^+^ and CD8^+^ SP cell populations (Fischer, Katayama, Pagès, Pouysségur, & Hedrick, 2005; Lesourne et al., 2009; Neilson, Winslow, Hur, & Crabtree, 2004; D. Wang et al., 2012). Therefore, we examined thymic positive selection in *Cic^f/f^;Vav1-Cre* mice. First, we analyzed thymocytes from 7-week-old WT, *Cic^f/f^;Cd4-Cre,* and *Cic^f/f^;Vav1-Cre* mice for the expression of CD69 and TCRβ using flow cytometry. The frequency of CD69^+^TCRβ^hi^ cells, which represent post-positive-selection thymocytes (Guo Fu et al., 2009), was significantly decreased in *Cic^f/f^;Vav1-Cre* mice compared with that of WT or *Cic^f/f^;Cd4-Cre* mice (Fig. 3A). This reduction was also observed in the thymus of *Cic^f/f^;Vav1-Cre* mice at 1 week of age (Fig. 3B). Next, we determined the effect of CIC deficiency on thymic positive selection in mice expressing the MHC class I-restricted H-Y TCR transgene specific to the H-Y male antigen (Kisielow, Blüthmann, Staerz, Steinmetz, & Von Boehmer, 1988) or the MHC class II-restricted OT-II TCR transgene specific to ovalbumin (Barnden, Allison, Heath, & Carbone, 1998) at 7–9 weeks of age. Defects in SP thymocyte development were much more severe in TCR-transgenic hematopoietic lineage cell-specific *Cic*-null (*H-Y;Cic^f/f^;Vav1-Cre* or *OT-II;Cic^f/f^;Vav1-Cre*) mice than in non-transgenic polyclonal *Cic^f/f^;Vav1-Cre* mice: CIC deficiency dramatically reduced the populations of CD8^+^ and CD4^+^ SP cells in female H-Y TCR transgenic and OT-II TCR transgenic mice, respectively (Figs. 3C and D, Fig. S2A). These results indicate that CIC controls the positive selection of thymocytes.

**Figure 3.**
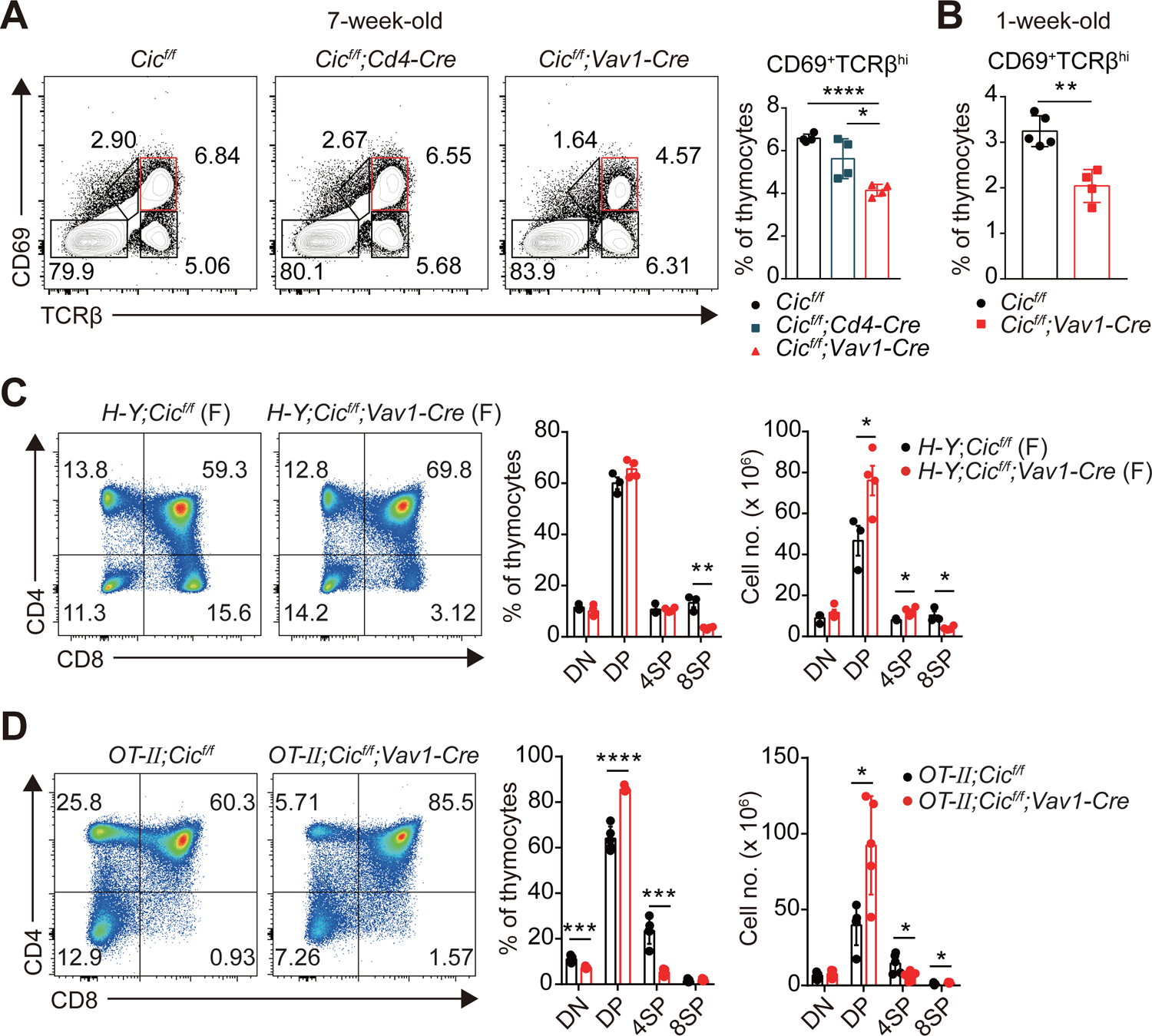
Defective positive selection in the absence of CIC. **(A)** Thymocytes from 7-week-old *Cic^f/f^*, *Cic^f/f^;Cd4-Cre*, and *Cic^f/f^;Vav1-Cre* mice were analyzed for surface expression of CD69 and TCRb. Representative flow cytometry plots (left) and the frequency of post-positive selection subset (CD69^+^TCRb^hi^) are shown (right). The CD69^+^TCRb^hi^ cell population is highlighted with a red box in the FACS plots. N = 4 for each group. **(B)** Frequencies of the post-positive selection subset (CD69^+^TCRb^hi^) of 1-week-old *Cic^f/f^* and *Cic^f/f^;Vav1-Cre* mice. N = 5 and 4 for *Cic^f/f^* and *Cic^f/f^;Vav1-Cre* mice, respectively. **(C)** Thymocytes from female *H-Y;Cic^f/f^* (N = 3) and *H-Y;Cic^f/f^*;*Vav1-Cre* (N = 4) mice were analyzed for CD4 and CD8 expression. Representative flow cytometry plots (left) and frequencies (middle) and numbers (right) of DN, DP, CD4^+^ SP (4SP), and CD8^+^ SP (8SP) cells are shown. **(D)** Thymocytes from *OT-II;Cic^f/f^* and *OT-II;Cic^f/f^;Vav1-Cre* mice were analyzed for CD4 and CD8 expression. Representative flow cytometry plots (left) and the frequencies (middle) and numbers (right) of DN, DP, CD4^+^ SP (4SP), and CD8^+^ SP (8SP) cells are shown. N = 5 for each group. The data are representative of at least two independent experiments. Bar graphs show data as means with SEM. *P < 0.05, **P < 0.01, ***P < 0.001, and ****P < 0.0001. The unpaired two-tailed Student’s t-test was used to calculate P values. See also Figure 3-source data 1.

### Impaired thymic negative selection in *Cic^f/f^;Vav1-Cre* mice

Most thymocytes expressing autoreactive TCRs are eliminated through negative selection during the DP and SP developmental stages (Klein et al., 2014). The strong TCR signal induced by the interaction of a self-reactive TCR with the self-pMHC molecule triggers apoptosis of autoreactive T cells (Hogquist & Jameson, 2014). Interestingly, we found that the expression of *Bcl2*, a representative anti-apoptotic gene (Vaux, Cory, & Adams, 1988) and an inhibitor of negative selection (Williams, Norton, Halligey, Kioussis, & Brady, 1998), was markedly increased in DP cells but not in DN and SP cells from *Cic^f/f^;Vav1-Cre* mice (Fig. 4A). Furthermore, the TCR stimulation-induced expression of Nur77, an orphan receptor that promotes apoptosis and negative selection of thymocytes (Cainan, Szychowski, Ka-Ming Chan, Cado, & Winoto, 1995), was significantly reduced in DP cells from *Cic^f/f^;Vav1-Cre* mice compared to that in WT mice (Fig. 4B). To determine whether CIC regulates thymic negative selection, we analyzed thymocytes from male *H-Y;Cic^f/f^* and *H-Y;Cic^f/f^;Vav1-Cre* mice by flow cytometry. As previously reported (Kisielow et al., 1988), massive negative selection of DP and CD8^+^ SP thymocytes was observed in male H-Y TCR transgenic mice (Fig. 4C), in which the male specific H-Y autoantigens are expressed in thymic antigen-presenting cells. The CD8^+^ SP population and total thymic cell number significantly increased in male *H-Y;Cic^f/f^;Vav1-Cre* mice compared to male *H-Y;Cic^f/f^* mice (Figs. 4C and D, Fig. S2B), indicating thymic negative selection defects in the absence of CIC. Taken together, our data demonstrate that CIC is required for the negative selection mediated by TCR activation-induced apoptosis.

**Figure 4.**
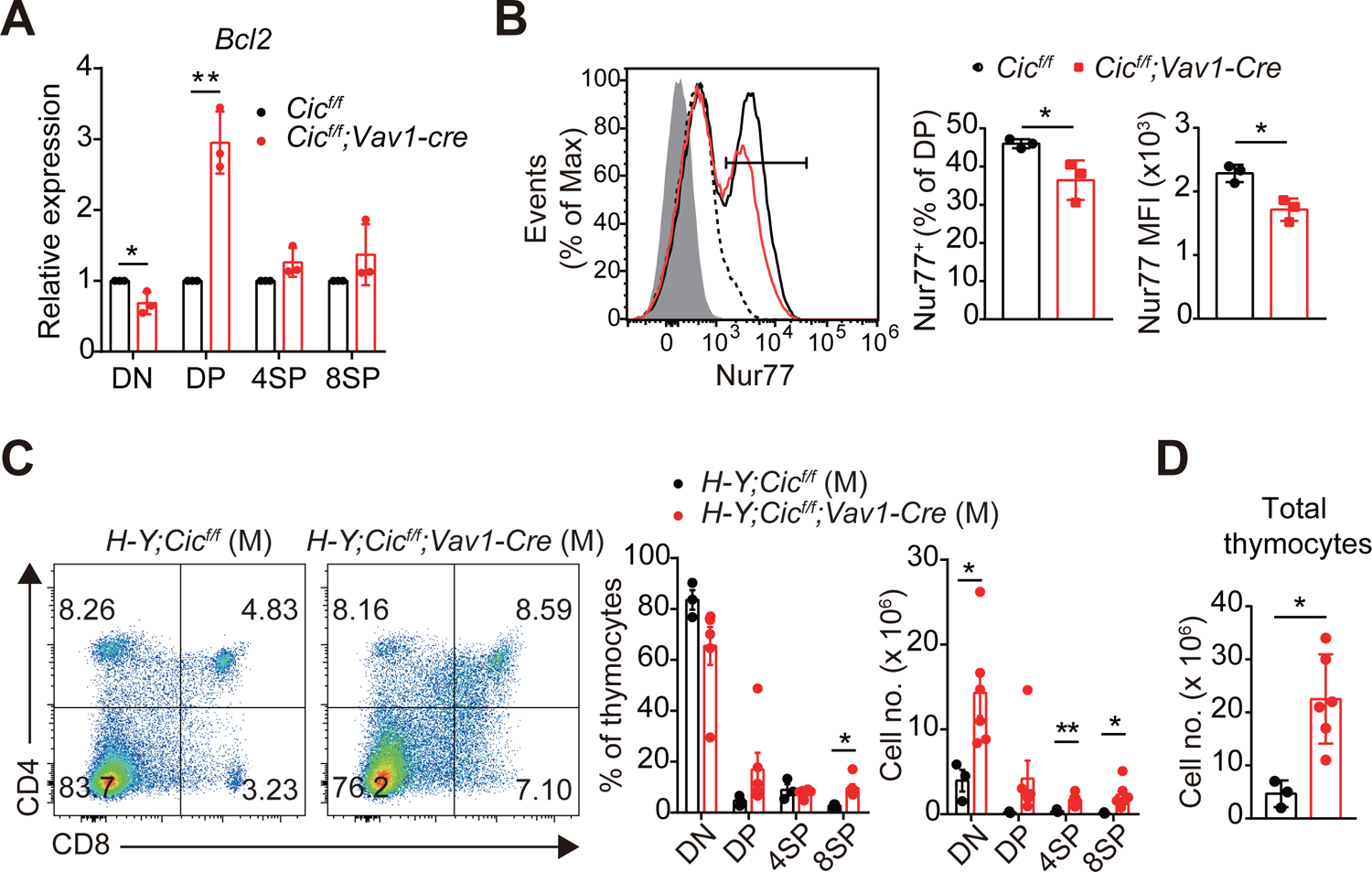
Defective negative selection in the absence of CIC. **(A)** qRT-PCR quantification of Bcl2 expression in DN, DP, CD4^+^ SP (4SP), and CD8^+^ SP (8SP) cells of *Cic^f/f^* and *Cic^f/f^;Vav1-Cre* mice. N = 3 for each group. **(B)** Flow cytometry of Nur77 expression in DP thymocytes from *Cic^f/f^* and *Cic^f/f^;Vav1-Cre* mice. Freshly isolated thymocytes were treated with plate-coated anti-CD3 (5 mg/ml) and anti-CD28 (10 mg/ml) for 2 h and then subjected to flow cytometry. Representative histograms for Nur77 expression in DP thymocytes of *Cic^f/f^* (black line) and *Cic^f/f^;Vav1-Cre* (red line) mice overlaid with isotype control (gray shaded) and unstimulated control (dotted line) histograms (left), the frequency of Nur77^+^ DP cells (middle), and Nur77 MFI of DP thymocytes (right) are presented. N = 3 for each genotype. Data are representative of two independent experiments. **(C and D)** Thymocytes from male *H-Y;Cic^f/f^* (N = 3) and *H-Y;Cic^f/f^;Vav1-Cre* (N = 6) mice were analyzed for CD4 and CD8 expression. **(C)** Representative flow cytometry plots (left) and frequencies (middle) and numbers (right) of DN, DP, CD4^+^ SP (4SP), and CD8^+^ SP (8SP) cells are shown. **(D)** Total thymocyte numbers. Data are representative of two independent experiments. Bar graphs show data as means with SEM. *P < 0.05 and **P < 0.01. The unpaired two-tailed Student’s t-test was used to calculate P values. See also Figure 4-source data 1.

### Changes in TCR repertoire of CD4^+^ SP thymocytes due to CIC deficiency

Developing T cells mature into SP cells through thymic selection processes; consequently, numerous T cells with various TCRs constitute the TCR repertoire. Because positive and negative selection determine the fate of T cells based on their TCR, defects in these processes can lead to changes in the TCR repertoire (J. Lu et al., 2019; Martínez-Riaño et al., 2019). To determine whether CIC deficiency alters the TCR repertoire of SP thymocytes, we prepared thymic regulatory T (Treg, CD4^+^CD25^+^GFP^+^) and non-Treg (CD4^+^CD25^-^GFP^-^) cells from *Foxp3-GFP;Cic^f/f^* and *Foxp3-GFP;Cic^f/f^;Vav1-cre* mice, and performed RNA sequencing for TCRα and TCRβ chain genes in each sample. The analysis of hypervariable complementarity-determining region 3 (CDR3) of TCRα and TCRβ chains revealed that in CIC-deficient cells, the CDR3 amino acid length distributions of TCR were generally longer than in WT cells, with some differences between Treg and non-Treg cells or between TCRα and TCRβ chains (Fig. S3A). This result suggests that CIC regulates the thymic selection process because pre-selection TCR chains in DN and DP thymocytes that have not undergone positive and negative selection show longer CDR3 amino acid length distributions than mature post-selection TCRs in CD44^-^CD62L^+^ naive T cells (J. Lu et al., 2019).

We also analyzed the usage of the variable (V) and joining (J) segments of the TCRα and TCRβ chains. The diversity of V and J segment combinations in the TCRβ chain was increased in CIC-deficient Treg cells compared to WT Treg cells, whereas it was comparable between WT and CIC-deficient non-Treg cells (Figs. S3B). Moreover, for both TCRα and TCRβ chains, the usage frequency of multiple V and J segments was different between WT and CIC-deficient Treg cells, whereas there were no significant differences in the non-Treg cell population (Figs. S3C-F). Taken together, these data suggest that impaired thymic selection due to CIC deficiency generated an abnormal Treg cell population with a distinct TCR repertoire in the thymus.

### Attenuated TCR signaling in CIC-deficient DP thymocytes

Because both positive and negative selection are mediated by TCR signaling strength (Klein et al., 2014), we investigated whether CIC regulates the TCR signaling pathway. First, we analyzed the surface expression of CD5, which has levels strongly correlated with TCR signaling intensity (Azzam et al., 1998), on DP and SP thymocytes from WT, *Cic^f/f^;Cd4-Cre,* and *Cic^f/f^;Vav1-Cre* mice. CD5 levels were substantially decreased in DP cells from *Cic^f/f^;Vav1-Cre* mice compared to those from WT or *Cic^f/f^;Cd4-Cre* mice, whereas this change was subtle in the CD4^+^ SP cell population and absent in the CD8^+^ SP cell population (Fig. 5A). This result was recapitulated in TCR transgenic mice (Figs. S2C and D). Next, we examined the activation of the TCR signaling pathway in thymocytes from WT, *Cic^f/f^;Cd4-Cre,* and *Cic^f/f^;Vav1-Cre* mice upon TCR stimulation. As with CD5 levels, calcium influx sharply decreased in DP cells from *Cic^f/f^;Vav1-Cre* mice, but not in SP thymocytes (Fig. 5B). We also observed a moderate decrease in calcium influx in DP cells from *Cic^f/f^;Cd4-Cre* mice (Fig. 5B), suggesting that CIC sensitively regulates TCR activation-induced calcium influx in DP thymocytes. To further evaluate CIC regulation of TCR signaling in DP thymocytes, we assessed the activation of key TCR signaling cascade components in DP cells of WT, *Cic^f/f^;Cd4-Cre,* and *Cic^f/f^;Vav1-Cre* mice after TCR stimulation by treatment with anti-CD3 and anti-CD4. Among the components tested, including ZAP-70, PLCγ, ERK, JNK, and p38, phospho-ERK levels were significantly decreased in DP cells from *Cic^f/f^;Vav1-Cre* mice compared to those from WT and *Cic^f/f^;Cd4-Cre* mice (Figs. 5C and D). Together, these results demonstrate that CIC deficiency attenuates TCR signaling by inhibiting calcium influx and ERK activation, especially in DP thymocytes.

**Figure 5.**
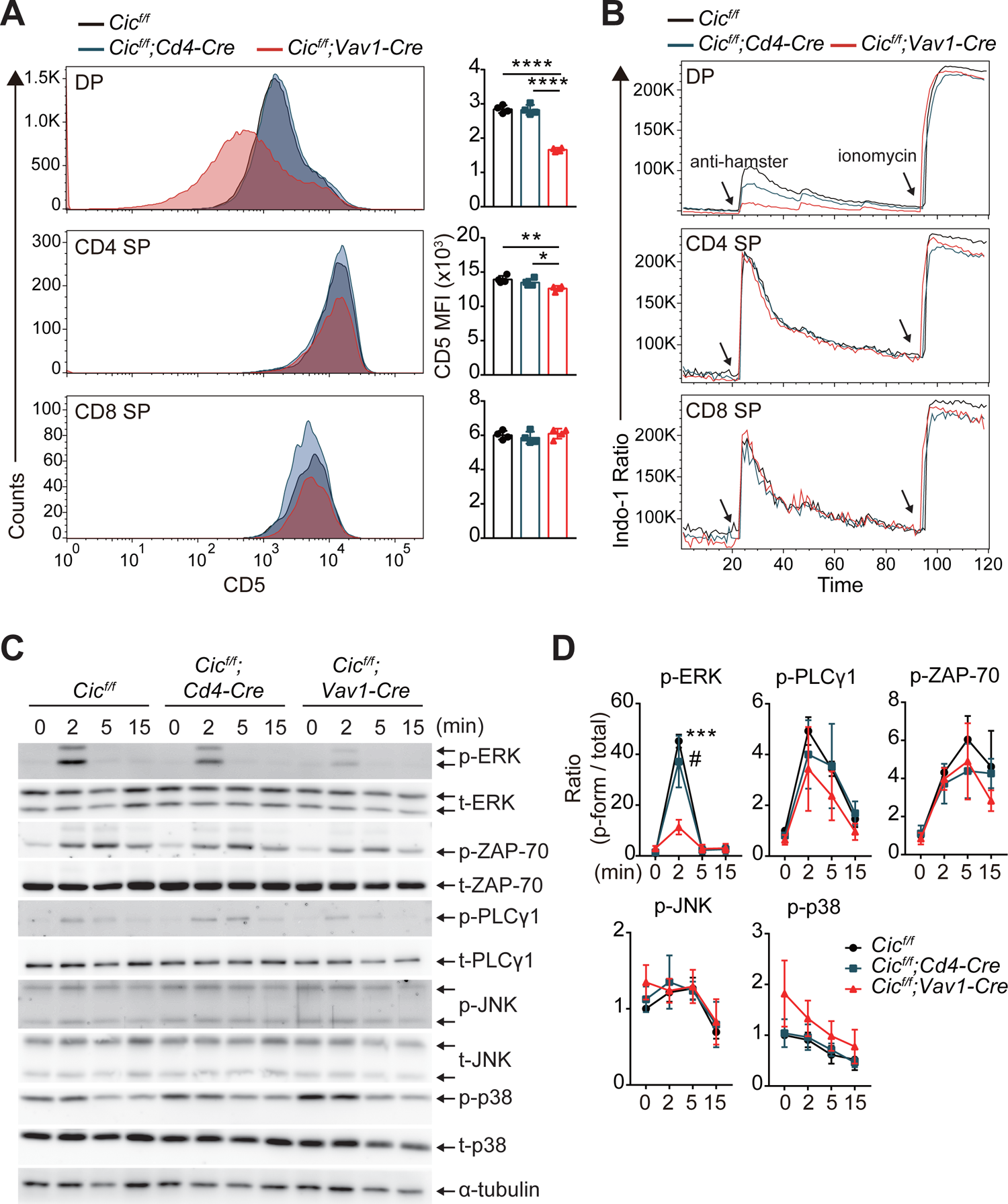
Attenuated TCR signaling in CIC-deficient DP thymocytes. **(A)** Thymocytes from 7-week-old *Cic^f/f^*, *Cic^f/f^;Cd4-Cre*, and *Cic^f/f^;Vav1-Cre* mice were FACS gated into the DP, CD4^+^ SP, and CD8^+^ SP cell population and CD5 MFI was measured. Representative histogram of CD5 expression in each cell population (left) and calculated CD5 MFI (right) are shown (N = 4 per group). **(B)** TCR stimulation-induced Ca2^+^ influx in DP, CD4^+^ SP, and CD8^+^ SP thymocytes from 7-week-old *Cic^f/f^*, *Cic^f/f^;Cd4-Cre*, and *Cic^f/f^;Vav1-Cre* mice. Data are representative of at least three independent experiments. **(C and D)** Western blot analysis for the activation of TCR cascade components in DP thymocytes from 7-week-old *Cic^f/f^*, *Cic^f/f^;Cd4-Cre*, and *Cic^f/f^;Vav1-Cre* mice. The sorted DP cells were stimulated with soluble anti-CD3 and anti-CD4 for indicated time points. **(C)** Representative western blot images. **(D)** Band densities were measured using ImageJ software and are presented as ratios of phosphorylated to total forms of each protein (N = 3). Graphs show data as means with SEM. *P < 0.05, **P < 0.01, ***P < 0.001, and ****P < 0.0001. #P< 0.05 (comparison between *Cic^f/f^;Cd4-Cre* and *Cic^f/f^;Vav1-Cre* mice). The unpaired two-tailed Student’s t-test was used to calculate P values. See also Figure 5-source data 1, 2.

### Abnormal thymic T cell development and reduced TCR signaling intensity in *Cic^f/f^;Vav1-Cre* mice are T cell-intrinsic

Although most of the cells that make up thymocytes are developing T cells, other types of immune cells, including B cells and dendritic cells, also exist in small proportions and participate in the regulation of T cell development (Klein et al., 2014). Because CIC expression was abolished in whole immune cells of *Cic^f/f^;Vav1-Cre* mice, it is unclear whether abnormal thymic T cell development and decreased TCR signaling in *Cic^f/f^;Vav1-Cre* mice were T cell-intrinsic or -extrinsic. To clarify this issue, we generated mixed BM chimeric mice by transferring the same number of BM cells from Thy1.1/Thy1.2 heterozygous WT and Thy1.1/Thy1.1 homozygous *Cic^f/f^;Vav1-Cre* mice into irradiated Thy1.2/Thy1.2 homozygous WT recipient mice (mixed WT:*Cic^f/f^;Vav1-Cre* BM chimera), and analyzed thymic T cells in the chimeras after 8 weeks of reconstitution (Fig. 6A). Similar to the observations in *Cic^f/f^;Vav1-Cre* mice (Figs. 1C and D), the frequency of DN4 and CD4^+^ SP cells was significantly decreased in the CIC-deficient cell compartment compared to the WT cell counterpart in the same mixed WT:*Cic^f/f^;Vav1-Cre* BM chimeric mice (Figs. 6B and C). The frequency of CD69^+^TCRβ^hi^ cells was also significantly lower in the CIC-deficient cell compartment relative to the WT cell compartment (Fig. 6D), indicating that the impaired positive selection in *Cic^f/f^;Vav1-Cre* mice was T cell-intrinsic. Furthermore, similar to the results in Fig. 5A, CD5 levels were most strongly reduced in DP thymocytes derived from *Cic^f/f^;Vav1-Cre* BM cells (Fig. 6E). Of note, the frequency of CD69^+^TCRβ^hi^ cells and CD5 levels in DP thymocytes were also significantly decreased in *Cic^f/f^;pLck-Cre* mice, another T cell-specific *Cic* null mice, compared to that in WT mice; however, the fold decrease was more dramatic in *Cic^f/f^;Vav1-Cre* mice than in *Cic^f/f^;pLck-Cre* mice (Figs.S4A and B). Although *pLck-Cre* is expressed during the DN2 cell stage (P. P. Lee et al., 2001), a small amount of CIC still existed in DP thymocytes from *Cic^f/f^;pLck-Cre* mice (Fig. S4C), explaining why *Cic^f/f^;pLck-Cre* mice showed a less dramatic decrease in the frequency of CD69^+^TCRβ^hi^ cells and CD5 levels in DP thymocytes compared to *Cic^f/f^;Vav1-Cre* mice. Overall, these data suggest that defects in thymic T cell development and TCR signaling in *Cic^f/f^;Vav1-Cre* mice are T cell-intrinsic.

**Figure 6.**
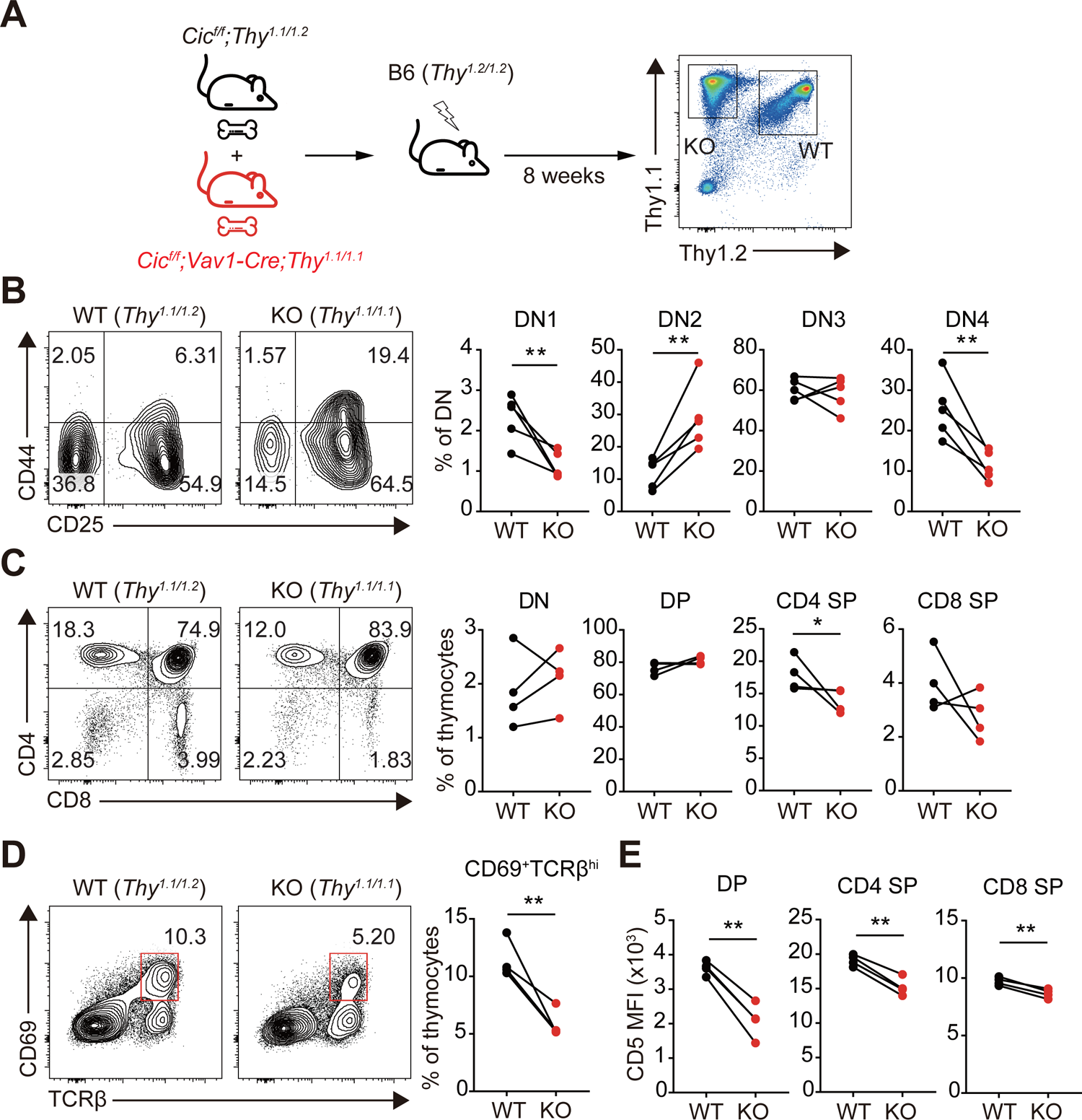
Altered T cell development and TCR intensity in *Cic^f/f^;Vav1-Cre* mice are caused by CIC loss in T cells. **(A)** Schematic for the generation and analysis of mixed BM chimeric mice. Equal numbers of BM cells from *Cic^f/f^;Thy^1.1/1.2^* (WT) and *Cic^f/f^;Vav1-Cre;Thy^1.1/1.1^* (KO) were mixed and transferred to irradiated B6 (*Thy^1.2/1.2^*) recipient mice (N = 4). A representative FACS plot image showing the thymocytes of each origin is presented. **(B-D)** Flow cytometry of thymocytes from the mixed BM chimeras for the frequencies of DN subsets based on CD44 and CD25 expression **(B)**, DN, DP, CD4^+^ SP, and CD8^+^ SP cells **(C)**, and post-positive selection subset (CD69^+^TCRb^hi^). The CD69^+^TCRb^hi^ cell population is highlighted with a red box in the flow cytometry plots **(D)**. **(E)** Flow cytometry of surface expression levels of CD5 in DP, CD4^+^ SP, and CD8^+^ SP thymocytes derived from WT and KO BM cells in the same BM chimeric mice. Data are representative of two independent experiments. Graphs show data as means with SEM. *P < 0.05, **P < 0.01, and ***P < 0.001. The unpaired two-tailed Student’s t-test was used to calculate P values. See also Figure 6-source data 1.

### Identification of CIC target genes responsible for attenuated TCR signaling in CIC-deficient DP thymocytes

To understand how CIC regulates the thymic selection process and TCR signaling in DP cells at the molecular level, we analyzed gene expression profiles of DP thymocytes from WT and *Cic^f/f^;Vav1-cre* mice by RNA sequencing. A total of 482 differentially expressed genes (DEGs; fold change > 2 and P-value < 0.05), including 263 upregulated and 219 downregulated genes, were identified in CIC-deficient DP cells (Table S1). Gene Ontology (GO) analysis revealed that genes involved in the inactivation of MAPKs and anti-apoptotic processes were significantly enriched among the upregulated DEGs (Table S2), consistent with the characteristics found in CIC-deficient DP thymocytes. We also identified several known CIC target genes among the DEGs, including *Etv1*, *Etv4*, *Etv5*, *Spry4*, *Dusp6*, *Dusp4,* and *Spred1* (Fryer et al., 2011; Weissmann et al., 2018; Yang et al., 2017) (Fig. 7A and Table S1). Of these, *Spry4*, *Dusp4*, *Dusp6,* and *Spred1* were of particular interest because they are negative regulators of ERK activation (Kidger & Keyse, 2016; Sasaki et al., 2003; Wakioka et al., 2001). Murine SPRY4 suppresses Ras-independent ERK activation by binding to Raf1 (Sasaki et al., 2003), whereas it represses insulin receptor and EGFR-induced ERK signaling upstream of Ras in humans (Leeksma et al., 2002). SPRED1 also inhibits the activation of ERK by suppressing Raf activation (Wakioka et al., 2001). DUSP6 specifically dephosphorylates activated ERK1/2, whereas DUSP4 acts on ERK as well as other MAPKs, such as JNK and p-38 (Caunt & Keyse, 2013). Additionally, SPRY4 suppresses calcium mobilization in HEK293T cells by inhibiting phosphatidylinositol 4,5-biphosphate (PIP_2_) hydrolysis induced by VEGF-A (Ayada et al., 2009). Thus, we investigated the association between derepression of *Spry4*, *Dusp4*, *Dusp6,* and *Spred1* and attenuated TCR signaling in CIC-deficient DP thymocytes. First, we measured the levels of *Spry4*, *Dusp4*, *Dusp6,* and *Spred1* in DN, DP, CD4^+^ SP, and CD8^+^ SP thymocytes from WT and *Cic^f/f^;Vav1-Cre* mice by qRT-PCR. The expression of all four genes was significantly upregulated in DP cells from *Cic^f/f^;Vav1-Cre* mice relative to those from WT mice (Fig. 7B), verifying the RNA sequencing data (Table S1 and Fig. 7A). The expression of all four genes was also upregulated in CD4^+^ and CD8^+^ SP thymocytes from *Cic^f/f^;Vav1-Cre* mice; *Spry4* and *Dusp6* were more dramatically upregulated in DP cells than in SP cells (Fig. 7B). Considering the different gene expression changes in each cell type, as well as the previously known functional significance in the regulation of ERK activation and calcium influx, two of the four CIC target genes, *Spry4* and *Dusp6*, were selected for further investigation of their effect on TCR signaling in DP thymocytes.

**Figure 7.**
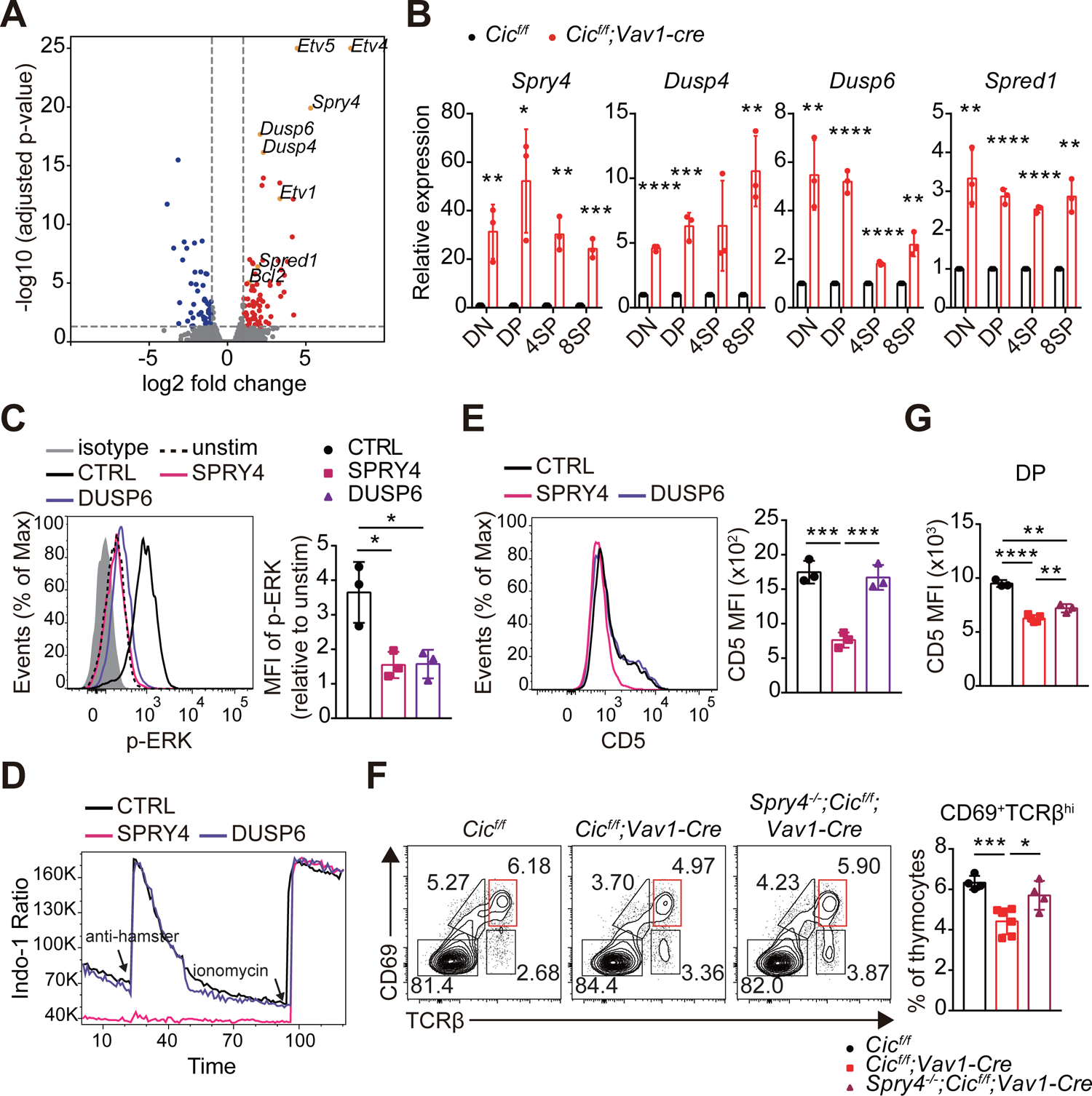
Identification of CIC target genes regulating TCR signaling in DP thymocytes. **(A)** Volcano plot showing the DEGs in CIC-deficient DP thymocytes (fold change >2 and adjusted P-value < 0.05). CIC target genes and *Bcl2* are marked at each corresponding dot. **(B)** qRT-PCR quantification of expression of *Spry4*, *Dusp4*, *Dusp6*, and *Spred1* in DN, DP, CD4^+^ SP (4SP), and CD8^+^ SP (8SP) thymocytes from *Cic^f/f^* and *Cic^f/f^;Vav1-Cre* mice. N = 3 for each group. **(C-E)** Effects of overexpression of SPRY4 and DUSP6 on TCR signaling in DP cells. Thymocytes were infected with retroviruses expressing either *Spry4* or *Dusp6* and subjected to flow cytometry for ERK activation **(C)**, Ca2^+^ influx **(D)**, and CD5 expression **(E)** in DP thymocytes. Three independent experiments were performed. **(F)** Thymocytes from 7-week-old *Cic^f/f^*, *Cic^f/f^;Vav1-Cre*, and *Spry4^-/-^;Cic^f/f^;Vav1-Cre* mice were analyzed for the surface expression of CD69 and TCRb. Representative FACS plots (left) and the frequency of CD69^+^TCRb^hi^ cells (right) are shown. The CD69^+^TCRb^hi^ cell population is highlighted with a red box in the FACS plots. N = 4, 6, and 4 for *Cic^f/f^*, *Cic^f/f^;Vav1-Cre*, and *Spry4^-/-^;Cic^f/f^;Vav1-Cre* mice, respectively. **(G)** CD5 levels in DP thymocytes from the mice used in **(F)**. N = 3, 5, and 3 for *Cic^f/f^*, *Cic^f/f^;Vav1-Cre*, and *Spry4^-/-^;Cic^f/f^;Vav1-Cre* mice, respectively. Bar graphs show data as means with SEM. *P < 0.05, **P < 0.01, ***P < 0.001, and ****P < 0.0001. The unpaired two-tailed Student’s t-test was used to calculate P values. See also Figure 7-source data 1.

We infected thymocytes with retroviruses expressing *Spry4* or *Dusp6* and analyzed phospho-ERK and CD5 levels and calcium influx by flow cytometry. The levels of phospho-ERK were greatly decreased in both SRPY4*-* and DUSP6-overexpressing DP thymocytes treated with anti-CD3 (Fig. 7C), confirming their inhibitory effect on ERK activation (Groom, Sneddon, Alessi, Dowd, & Keyse, 1996; Muda et al., 1996; Sasaki et al., 2003). In contrast, calcium influx induced by TCR stimulation was almost completely suppressed in DP thymocytes by SPRY4 overexpression but not by DUSP6 overexpression (Fig. 7D). Moreover, CD5 expression was substantially reduced in SRPY4*-*overexpressing cells, but not in DUSP6*-*overexpressing cells (Fig. 7E). These data imply that *Spry4* derepression might critically contribute to attenuated TCR signaling in CIC-deficient DP thymocytes.

Finally, to determine whether impaired thymic selection and TCR signaling in *Cic^f/f^;Vav1-Cre* mice were caused by derepression of *Spry4*, we generated *Cic* and *Spry4* double mutant (*Spry4^-/-^;Cic^f/f^;Vav1-Cre*) mice and analyzed thymic T cells in WT, *Cic^f/f^;Vav1-Cre,* and *Spry4^-/-^;Cic^f/f^;Vav1-Cre* mice at 7–8 weeks of age. The decreased frequency of CD69^+^TCRβ^hi^ cells and CD5 levels of DP thymocytes in *Cic^f/f^;Vav1-Cre* mice were significantly, but only partially, rescued in *Spry4^-/-^;Cic^f/f^;Vav1-Cre* mice (Figs. 7F and G), suggesting a partial recovery of the CIC deficiency-mediated defective positive selection process by loss of SPRY4. However, the decreased DN4 cell frequency and Nur77 expression in DP thymocytes were not rescued in *Spry4^-/-^;Cic^f/f^;Vav1-Cre* mice (Figs. S5A and B), indicating that *Spry4* derepression alone is not sufficient to induce abnormal DN cell development in *Cic^f/f^;Vav1-Cre* mice and suppress TCR activation-induced apoptosis in CIC-deficient DP thymocytes. Taken together, our findings demonstrate that CIC tightly controls thymic T cell development and TCR signaling by repressing multiple CIC target genes, including *Dusp4*, *Dusp6*, *Spred1*, and *Spry4*, in a cell type- and/or developmental stage-specific manner.

## DISCUSSION

Our study uncovered the role of CIC in thymic T cell development via in-depth analyses of thymocytes in various CIC-deficient mouse models. CIC deficiency decreased the frequency of DN4 thymocytes that underwent β-selection mediated by pre-TCR and Notch signaling (Dose et al., 2006; Kreslavsky et al., 2012). However, the decreased DN4 cell population did not prevent the formation of thymic DP cells in *Cic^f/f^;Vav1-Cre* mice. Thus, CIC deficiency could impair the β-selection process; nonetheless, this defect was not sufficient to significantly affect the transition from the DN to DP stage of thymic T cell development in *Cic^f/f^;Vav1-Cre* mice. Interestingly, similar to *Cic^f/f^;Vav1-Cre* mice, ERK-deficient mice also exhibit a partial block in DN3-to-DN4 maturation without defects in maturation to the DP stage of development (Fischer et al., 2005). Because TCR stimulation-induced ERK activation was markedly suppressed in CIC-deficient DP thymocytes (Figs. 5C and D), we inferred that formation of the abnormal DN subset in the thymus of *Cic^f/f^;Vav1-Cre* mice was, at least in part, caused by reduced ERK activity in DN cells deficient in CIC. Consistent with this inference, the *Spry4*, *Dusp4*, *Dusp6,* and *Spred1* genes, involved in the inhibition of ERK activation (Kidger & Keyse, 2016; Sasaki et al., 2003; Wakioka et al., 2001), were significantly derepressed in CIC-deficient DN cells (Fig. 7B). Further study on how CIC controls pre-TCR signaling is critical to better understand CIC regulation of thymic T cell development during the DN stage.

CIC deficiency, especially in DP thymocytes, disrupts the positive and negative selection of thymocytes, as evidenced by the impaired thymic selection process in *Cic^f/f^;Vav1-Cre* mice but not in *Cic^f/f^;Cd4-Cre* mice, which express significant amounts of CIC proteins in DP thymocytes. Moreover, CIC loss substantially suppressed TCR signaling in DP thymocytes but not in SP cells. Although *Spry4*, *Dusp4*, *Dusp6,* and *Spred1* were all significantly derepressed in SP and DP cells from *Cic^f/f^;Vav1-Cre* mice, among these four CIC target genes, *Spry4* and *Dusp6* were more dramatically derepressed in DP cells than in SP cells in the absence of CIC (Fig. 7B). These results suggest that *Spry4* and *Dusp6* may be CIC target genes primarily responsible for the CIC deficiency-mediated dysregulation of TCR signaling in DP cells and thymic selection processes. However, removal of the *Spry4* alleles only partially recovered the TCR signal intensity and positive selection, indicating that CIC regulates multiple target genes, including *Spry4*, to elaborately control TCR signaling and thymic T cell development. Notably, the hyperactivation of peripheral T cells and expansion of the Tfh cell population in *Cic^f/f^;Vav1-Cre* mice were also not rescued in *Spry4^-/-^;Cic^f/f^;Vav1-Cre* mice (Figs. S5C and D). These findings suggest that the partial recovery of the TCR signal intensity and positive selection process by loss of SPRY4 was insufficient to ameliorate the abnormal peripheral T cell phenotypes in *Cic^f/f^;Vav1-Cre* mice or that derepression of CIC target genes other than *Spry4* could lead to T cell hyperactivation and enhanced Tfh cell formation in *Cic^f/f^;Vav1-Cre* mice. Concordantly, we have shown that CIC deficiency promotes Tfh cell differentiation via *Etv5* derepression (S. Park et al., 2017). In contrast, CIC deficiency-induced thymic T cell phenotypes were not rescued in *Cic^f/f^;Etv5^f/f^;Vav1-Cre* mice (data not shown). Overall, these findings suggest that CIC regulates various target genes with differential effects to broadly control thymic T cell development and peripheral T cell activation and differentiation. Our sequencing analysis for *Tcra* and *Tcrb* mRNAs revealed that CIC deficiency substantially altered the TCR repertoire of thymic Treg cells. On the other hand, this alteration was not clearly detected in the non-Treg cell population. Based on the affinity model of thymocyte selection, thymic Treg cell differentiation occurs within a window between positive selection (low affinity of the TCR-self-pMHC interaction) and negative selection (high affinity of the TCR-self-pMHC interaction) (Klein et al., 2014). Due to impaired negative selection, some Treg cells could be derived from self-reactive T cells that were supposed to be eliminated by negative selection in *Cic^f/f^;Vav1-Cre* mice. Therefore, the changed TCR repertoire in CIC-deficient Treg cells might result from defective negative selection. The distribution of CDR3 amino acid length of the TCRβ chain was changed in CIC-deficient non-Treg cells (Fig. S3A). On the other hand, the diversity of V and J segment combinations in the TCRβ chain and the usage frequency of V and J segments were comparable between WT and CIC-deficient non-Treg cells (Figs. S3B-F). However, these data are limited in determining whether the TCR repertoire is normal in CIC-deficient non-Treg cells, since there is no information on the combination of TCRα and TCRβ chains at the single cell level. To clarify this issue, single cell RNA sequencing or TCR repertoire analysis using transgenic mice with fixed TCRα/TCRβ chain should be conducted.

This study provides insight into how CIC controls autoimmunity. Because defects in thymic selection lead to the breakdown of central tolerance (Xing & Hogquist, 2012), it is conceivable that CIC deficiency during thymic T cell development generates autoreactive T cells, inducing autoimmunity. Therefore, CIC potentially suppresses autoimmunity by controlling the thymic selection process, as well as Tfh cell differentiation (S. Park et al., 2017). The more severe autoimmune-like phenotypes in *Cic^f/f^;Vav1-Cre* mice than in *Cic^f/f^;Cd4-Cre* mice (S. Park et al., 2017) highlight the significant contribution of the impaired thymic selection process to CIC deficiency-induced autoimmunity. To clarify the importance of each regulatory step in the suppression of autoimmunity, it will be necessary to better understand the molecular mechanisms underlying CIC regulation of thymic T cell development and Tfh cell differentiation. Additionally, studies on the role of CIC in T cell development and Tfh cell differentiation in humans should be conducted in the future to improve our understanding of the pathogenesis of autoimmune diseases, such as systemic lupus erythematosus.

## MATERIALS AND METHODS

### Mice

All mice were maintained on a C57BL/6 background. *Cic-*floxed (H. C. Lu et al., 2017; S. Park et al., 2017), *Vav1-Cre* (de Boer et al., 2003), *CD4-Cre* (P. P. Lee et al., 2001), *pLck-Cre* (P. P. Lee et al., 2001), FLAG-tagged *Cic* knock-in (*Cic^FLAG/FLAG^*) (S. Park et al., 2019), H-Y TCR transgenic (Kisielow et al., 1988), OT-II TCR transgenic (Barnden et al., 1998), and *Foxp3^EGFP^* (Lin et al., 2007) mice have been described previously. *Spry4^-/-^* mice were generated using *Spry4^tm1a(KOMP)Mbp^* embryonic stem cells obtained from the UC Davis KOMP repository. All experiments were conducted with age-matched mice, and 7–9 week-old mice were used unless otherwise indicated. Mice from both sexes were randomly allocated to experimental groups. All mice were maintained in a specific pathogen-free animal facility under a standard 12 h light/12 h dark cycle. Mice were fed standard rodent chow and provided with water *ad libitum*. All experiments were approved by the Institutional Animal Care and Use Committee of Pohang University of Science and Technology.

### Cell line

The Platinum-E (Plat-E) retroviral packaging cell line (Cell Biolabs) was grown in Dulbecco’s modified Eagle’s medium (DMEM, Welgene) supplemented with 10% fetal bovine serum (FBS, Welgene), and penicillin/streptomycin (Gibco). Cells were cultured in a 37°C incubator with 5% CO_2_. Mycoplasma contamination was routinely tested using the e-Myco plus Mycoplasma detection kit (INtRON Bio).

### Flow cytometry

Surface staining was performed using the following fluorescence-labeled antibodies against: CD4 (RM4-5; Biolegend, BD Biosciences), CD5 (53-7.3; Biolegend, eBioscience), CD8α (53-6.7; Biolegend), CD11b (M1/70; Biolegend), CD11c (N418, Biolegend), CD19 (1D3; BD Biosciences), CD25 (PC61.5; Tonbo Biosciences), CD44 (IM7, BD Biosciences), CD62L (MEL-14; BD Biosciences), CD69 (H1.2F3; Biolegend), CD90.1 (HIS51; eBioscience), CD90.2 (53-2.1; Biolegend), CXCR5 (2G8; BD Biosciences), TCRβ (H57-597; Tonbo Biosciences), TCRγ/δ (GL-3; eBioscience), NK-1.1 (PK136; Biolegend), TER-119 (TER-119; eBioscience), Gr-1 (RB6-8C5; eBioscience), TCR H-Y (T3.70; eBioscience), and PD-1 (RMP1-30; eBioscience). For CXCR5 staining, cells were incubated with biotinylated anti-CXCR5 for 30 min and then sequentially incubated with APC- or PerCP-Cy5.5-labelled streptavidin (eBioscience) with other surface antibodies. For Live/dead staining was performed using Fixable Viability Dye (FVD) eFluor 780 (eBioscience) or Ghost Dye Violet 510 (Tonbo Biosciences). Intracellular staining was performed using the Foxp3 staining buffer set (eBioscience). For FLAG-CIC and Nurr77 staining, fluorochrome-labeled antibodies to FLAG (L5; Biolegend) and Nur77 (12.14; eBioscience) were used. For phospho-ERK staining, cells were stained with FVD, fixed with BD Cytofix, and permeabilized with cold methanol. Permeabilized cells were stained with the p-ERK antibody (Cell Signaling Technology), and then fluorochrome-labeled antibodies to surface markers and secondary antibody against rabbit IgG (Biolegend) were added. The stained cells were analyzed using an LSRII Fortessa flow cytometer (BD Biosciences) or CytoFLEX LX (Beckman Coulter). Data were analyzed using FlowJo software (TreeStar).

### Cell sorting

Single-cell suspensions of thymocytes were stained for surface markers including lineage (TCRγ/δ, NK-1.1, TER-119, Gr-1, CD11b, CD11c, CD19) and then lin^-^ DN (CD4^-^CD8^-^), ISP (CD4^-^CD8^+^TCRβ^lo^CD24^hi^), DP (CD4^+^CD8^+^TCRβ^lo^), CD4^+^ SP (CD4^+^CD8^-^TCRβ^hi^), and CD8^+^ SP (CD4^-^CD8^+^TCRβ^hi^) cells were sorted. To sort DN3 (lin^-^CD4^-^CD8^-^CD44^lo^CD25^hi^) and DN4 (lin^-^CD4^-^CD8^-^CD44^lo^CD25^lo^) cells, CD8^-^ thymocytes were obtained by negative selection using the EasySep Mouse Streptavidin Rapid Spheres Isolation Kit (Stem Cell Technologies), then subjected to cell sorting. A MoFlo-XDP (Beckman Coulter) was used for cell sorting.

### Generation of BM chimeric mice

To create a mixed BM chimera, 1 × 10^6^ BM cells from each donor were mixed and injected intravenously into C57BL/6 recipient mice that had been irradiated (10 Gy). After 8 weeks of recovery, the mice were sacrificed; thymi were harvested and homogenized to prepare single-cell suspensions. Cells were then stained for flow cytometry.

### *A*nalysis of Nurr77 expression

To measure Nur77 expression, freshly isolated thymocytes were incubated with plate-coated anti-CD3 (5 μg/ml) and anti-CD28 (10 μg/ml) for 2 h, followed by staining of surface markers (CD4, CD8, and TCRβ) and intracellular staining of Nur77. Samples were analyzed using an LSRII Fortessa flow cytometer or CytoFLEX LX. Data were analyzed using FlowJo software.

### *In vitro* TCR stimulation

Total thymocytes or sorted DP cells were rested for 30 min in Roswell Park Memorial Institute (RPMI) medium (Welgene) at 37°C. Cells were then washed with T-cell medium (TCM, RPMI supplemented with 10% FBS [Welgene], 1% penicillin/streptomycin [Gibco], and 0.1% β-mercaptoethanol [Gibco]) and incubated with biotin-conjugated anti-CD3 (60 μg/ml, 145-2C11) and anti-CD4 (60 μg/ml, GK1.5) for 20 min on ice. Cells were washed and incubated for 5 min in ice with streptavidin (60 μg/ml, SouthernBiotech) for cross-linking and incubated at 37°C. To analyze p-ERK levels, unconjugated anti-CD3 (10 μg/ml, 145-2C11) and goat anti-hamster IgG (25 μg/ml, Jackson Immunoresearch) were used to stimulate TCR. One milliliter of cold PBS was added at the end of stimulation and cell pellets were lysed for western blotting or further stained with anti-p-ERK antibody for flow cytometry.

### Calcium influx measurement

Freshly isolated thymocytes or virus-transduced thymocytes were incubated with 4 μM Indo-1-AM (Invitrogen) in TCM at 37°C for 40 min. Cells were washed twice with TCM and incubated with soluble anti-CD3 (10 μg/ml) and fluorochrome-conjugated antibodies for surface markers (CD4, CD8, and TCRβ) in TCM on ice for 20 min. Cells were then washed with TCM and warmed before cross-linking. Goat anti-hamster IgG (25 μg/ml) was added to cross-link anti-CD3 and the signals were measured by flow cytometry. Ionomycin was added to ensure that T cells were effectively loaded with Indo-1. The emission wavelength ratios of Ca^2+^-bound to unbound Indo-1 were analyzed using an LSRII Fortessa flow cytometer.

### Plasmids and retroviral transduction

The coding sequences (CDSs) of mouse *Spry4* or *Dusp6* with enzyme sites XhoI/EcoRI was amplified by PCR and cloned into the MigR1 retroviral vector (Pear et al., 1998). Virus was generated through transient cotransfection of the Plat-E cells with the cloned retroviral vectors and pCL-Eco helper plasmid (Imgenex) (Naviaux, Costanzi, Haas, & Verma, 1996). Briefly, 2.5 × 10^6^ Plat-E cells were plated in 100-mm plates. The next day, cells were transfected with 3.6 μg of retroviral vector and 2.4 μg of pCL-Eco using FuGENE HD transfection reagent (Promega). Retrovirus-containing supernatants were harvested twice at 48 h and 72 h after transfection and filtered with a 0.22 μm syringe filter (Millipore). For retroviral transduction, 10^7^ thymocytes were mixed with 0.5 ml of the viral supernatant and 1.5 ml of fresh TCM in the presence of 4 μg/ml polybrene (Sigma) and spin-infected at 1000 *g* for 90 min at 25°C.

### Western blotting

Sorted cells or stimulated cells were lysed in RIPA buffer (50mM Tris-HCl pH 7.4, 150mM NaCl, 1mM PMSF, 1% NP-40, 0.5% sodium deoxycholate, 0.1% SDS, 1x Roche Complete Protease Inhibitor Cocktail, and 1x Roche Phosphatase Inhibitor Cocktail). Protein concentrations were measured using a BCA kit (Pierce). Equal amounts of protein samples were separated by 9% SDS-PAGE gel and transferred onto nitrocellulose membranes (Bio-Rad). The following primary antibodies were used: anti-CIC (homemade) (Kim et al., 2015), anti-PLCγ1 (#5690, Cell Signaling), anti-p-PLCγ1 (#2821, Cell Signaling), anti-ZAP-70 (#2705, Cell Signaling), anti-p-ZAP-70 (#2701, Cell Signaling Technology), anti-JNK (#9252, Cell Signaling), anti-p-JNK (#9251, Cell Signaling), anti-ERK (#9102, Cell Signaling), anti-p-ERK (#4370, Cell Signaling), anti-p38 (#9212, Cell Signaling), anti-p-p38 (#9211, Cell Signaling), anti-β-actin (#sc-47778, Santa Cruz), and anti-α-tubulin (#sc-398103, Santa Cruz). Membranes were incubated with secondary antibodies conjugated to horseradish peroxidase (HRP) and developed using Clarity Western ECL Substrate (Bio-Rad) or SuperSignal West Dura (Thermo Scientific). Images were acquired using an ImageQuant LAS 500 instrument (GE Healthcare).

### RNA isolation, cDNA synthesis, and qRT-PCR

Total RNA was extracted from sorted cells with RiboEx (GeneAll) and 0.2–1 μg was subjected to cDNA synthesis using the GoScript Reverse Transcription system (Promega) according to the manufacturer’s instructions. SYBR Green real-time PCR master mix (TOYOBO) was used for qRT-PCR analysis. Primers used for qRT-PCR are listed in Table S3.

### RNA sequencing and data analysis

Thymi from *Cic^f/f^* and *Cic^f/f^;Vav1-Cre* mice were dissected and homogenized into single-cell suspensions for cell sorting. DP thymocytes were sorted on the basis of surface markers of lin^-^CD4^+^CD8^+^TCRβ^lo^, and total RNA was extracted with RiboEx. The library for mRNA sequencing was generated using the TruSeq Stranded Total RNA LT Sample Prep Kit (Illumina) and sequencing was performed with NovaSeq 6000 (Illumina). Trimmed reads were mapped to the mouse reference genome (mm10 RefSeq) with HISAT2, and the transcripts were assembled using StringTie. DEGs were generated using edgeR and genes with fold changes > 2 and P-values < 0.05 were selected for Gene Ontology (GO) analyses on the basis of biological processes using the DAVID website (https://david.ncifcrf.gov/).

### TCR repertoire sequencing

Treg (CD4^+^CD8^-^CD25^+^GFP^+^) and non-Treg (CD4^+^CD8^-^GFP^-^) cells were prepared from *Foxp3-GFP;Cic^f/f^* and *Foxp3-GFP;Cic^f/f^;Vav1-cre* mice by cell sorting. Total RNA was extracted from 3-5 × 10^4^ Treg and 3 × 10^6^ non-Treg cells, then subjected to sequencing of mRNA encoding TCRα and TCRβ chains, performed by iRepertoire, Inc. (Huntsville, AL). Briefly, RNA was amplified using a commercially available multiplex primer mix covering the TCRα and TCRβ chains in two separate PCR reactions. Reverse transcription and subsequent PCR amplification (RT-PCR1) were performed using the Qiagen OneStep RT PCR mix (Qiagen). The cDNA was selected and unused primers were removed by SPRIselect bead selection (Beckman Coulter) followed by a second round of amplification performed with a pair of primers specific for communal sites engineered onto the 5ʹ end of the C- and V-primers used during RT-PCR1. The final constructed library included Illumina sequencing adapters and a 6-nucleotide internal barcode associated with the C-gene primer. After library preparation, paired-end sequencing was performed using the Illumina Miseq v3 600-cycle Reagent Kit (Illumina). The output of the immune receptor sequence covered the second framework region through the beginning of the constant region, including both CDR2 and CDR3. Raw sequencing data were analyzed using the previously described iRmap program (C. Wang et al., 2010).

## Data availability

The Gene Expression Omnibus (GEO) accession number for the RNA sequencing data of DP thymocytes reported in this paper is GSE173909. The GEO accession number for the TCR repertoire RNA sequencing is GSE173996. All data generated or analysed during this study are included in the manuscript and supporting files. Source data files have been provided for Figures 1, 2, 3, 4, 5, 6, and 7.

## Statistical analysis

Statistical analysis was performed using GraphPad Prism 7 (www.Graphpad.com, La Jolla, CA, USA). All experiments were independently performed more than three times. Student’s *t*-test (two-tailed, two-sample unequal variance) was used to obtain P-values between groups. Statistical significance was set at P < 0.05. Error bars indicate the standard error of the mean (SEM).

## ACKNOWLEDGMENTS

We thank Dr. Jaeho Cho and the Lee lab members for their helpful discussions and comments on this study. This work was supported by grants from the Samsung Science and Technology Foundation under project number SSTF-BA1502-14 and the National Research Foundation (NRF) of Korea (NRF-2021R1A2C3004006 and - 2017R1A5A1015366). JSP and JP were supported by the BK21 Program. HH was supported by a Global PhD Fellowship (NRF-2017H1A2A1042705).

## AUTHOR CONTRIBUTIONS

S.K. and Y.L. designed the study. S.K., G.Y.P., J.S.P., J.P., and H.H. performed the experiments. S.K. analyzed the data. S.K. and Y.L. wrote the manuscript.

---

**Supplemental Figure 1.**
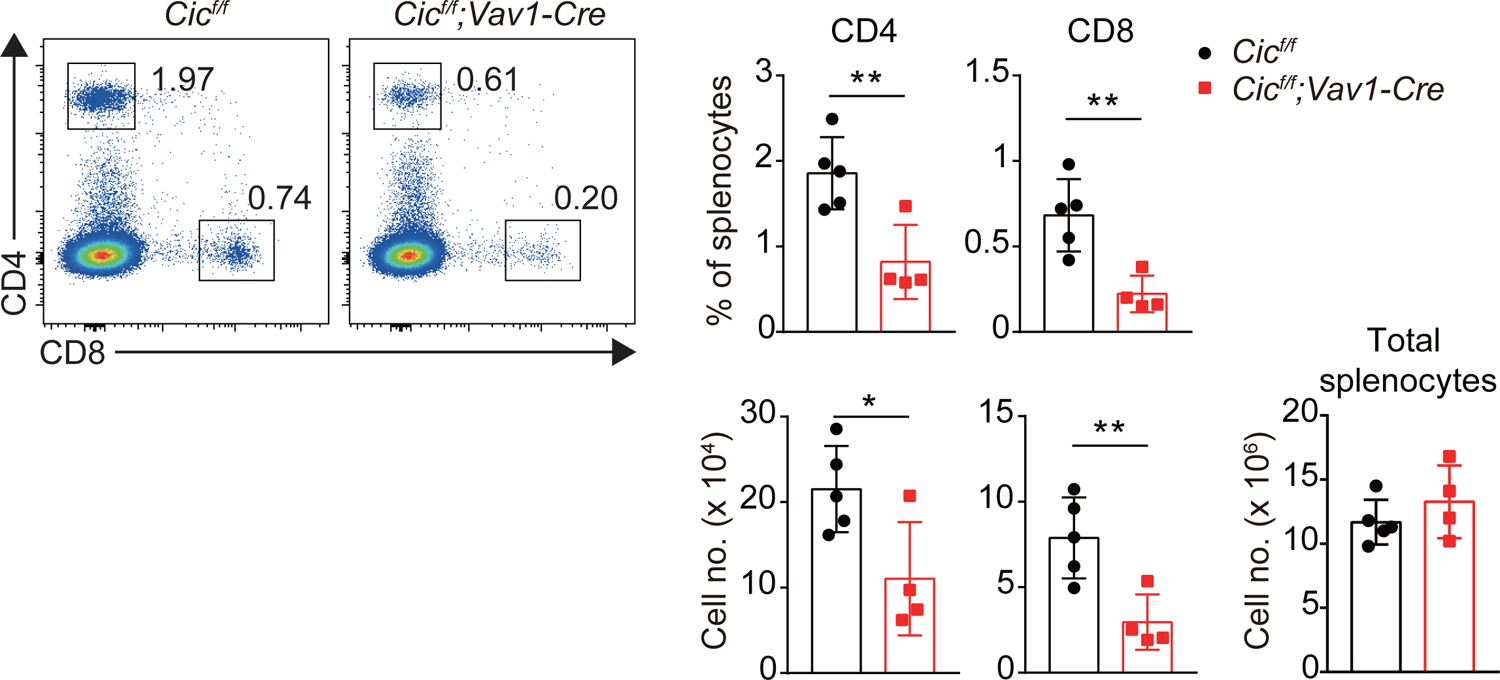
Decreased splenic T cell populations in *Cic^f/f^;Vav1-Cre* mice at 1 week of age. Flow cytometry of splenocytes from 1-week-old *Cic^f/f^* and *Cic^f/f^;Vav1-Cre* mice. Representative plot images of CD4 and CD8 expression in splenocytes, frequency and number of CD4^+^ and CD8^+^ T cells, and total splenocyte numbers are shown. Data represent two independent experiments. Bar graphs show data as means with SEM. *P < 0.05 and **P < 0.01.

**Supplemental Figure 2.**
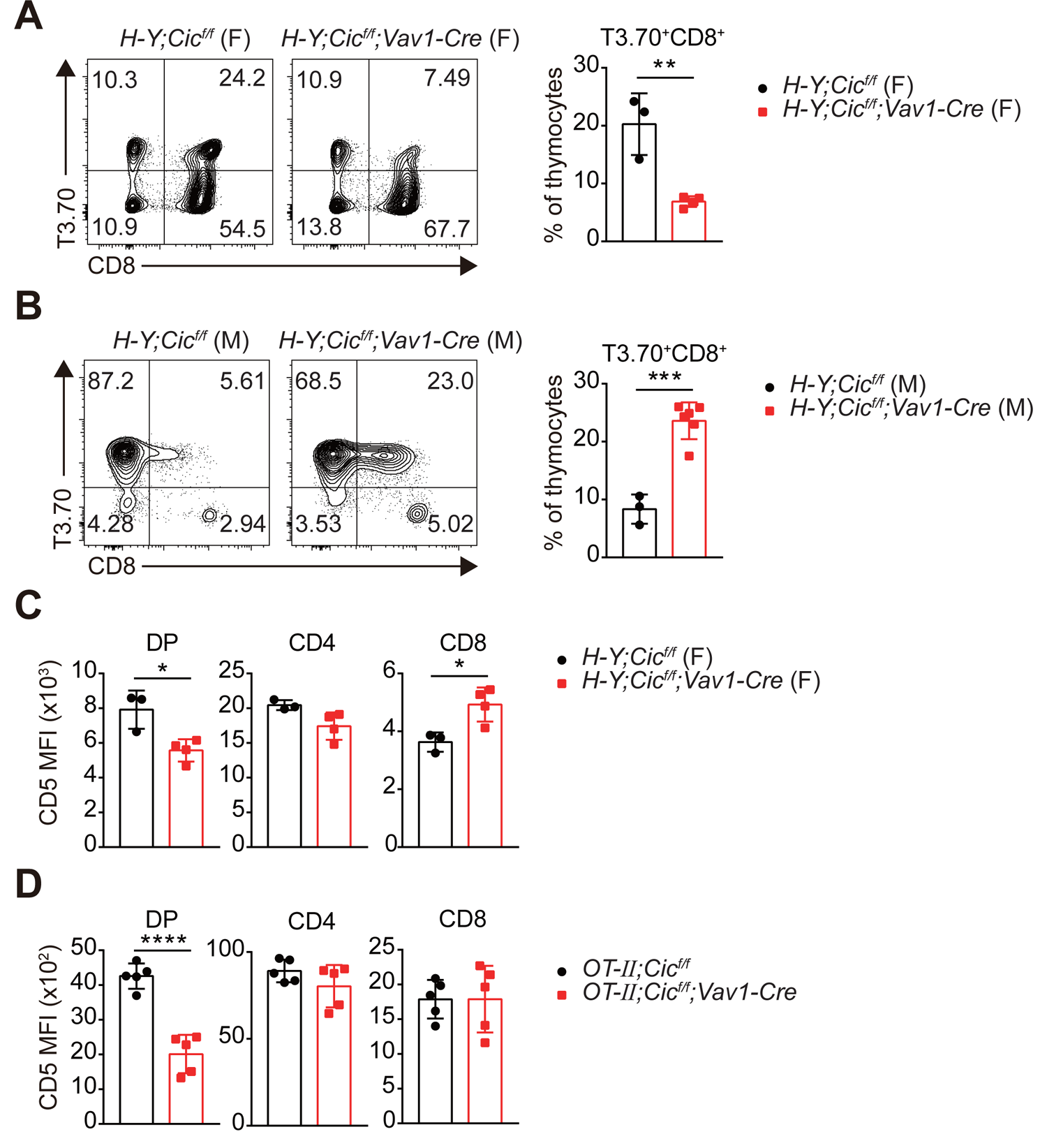
Analysis of H-Y TCR transgenic CD8^+^ T cells and CD5 levels in TCR transgenic thymocytes. **(A and B)** Flow cytometry of thymocytes from female and male *H-Y;Cic^f/f^* and *H-Y;Cic^f/f^*;*Vav1-Cre* mice for H-Y TCR (T3.70) and CD8 expression. Frequencies of H-Y TCR^+^ CD8^+^ cells among total thymocytes are presented. **(A)** Results for female mice. N = 3 and 4 for female *H-Y;Cic^f/f^* and *H-Y;Cic^f/f^*;*Vav1-Cre* mice, respectively. **(B)** Results for male mice. N = 3 and 6 for male *H-Y;Cic^f/f^* and *H-Y;Cic^f/f^*;*Vav1-Cre* mice, respectively. Data are representative of two independent experiments. **(C and D)** Thymocytes from 7-week-old WT and CIC-deficient TCR transgenic mice were FACS gated into the DP, CD4^+^ SP, and CD8^+^ SP cell population and CD5 MFI was measured. **(C)** Results for female H-Y TCR transgenic mice. N = 3 and 4 for *H-Y;Cic^f/f^* and *H-Y;Cic^f/f^*;*Vav1-Cre* mice, respectively. **(D)** Results for OT-II TCR transgenic mice. N = 5 for each group. Bar graphs show data as means with SEM. *P < 0.05, **P < 0.01, ***P < 0.001, and ****P < 0.0001.

**Supplemental Figure 3.**
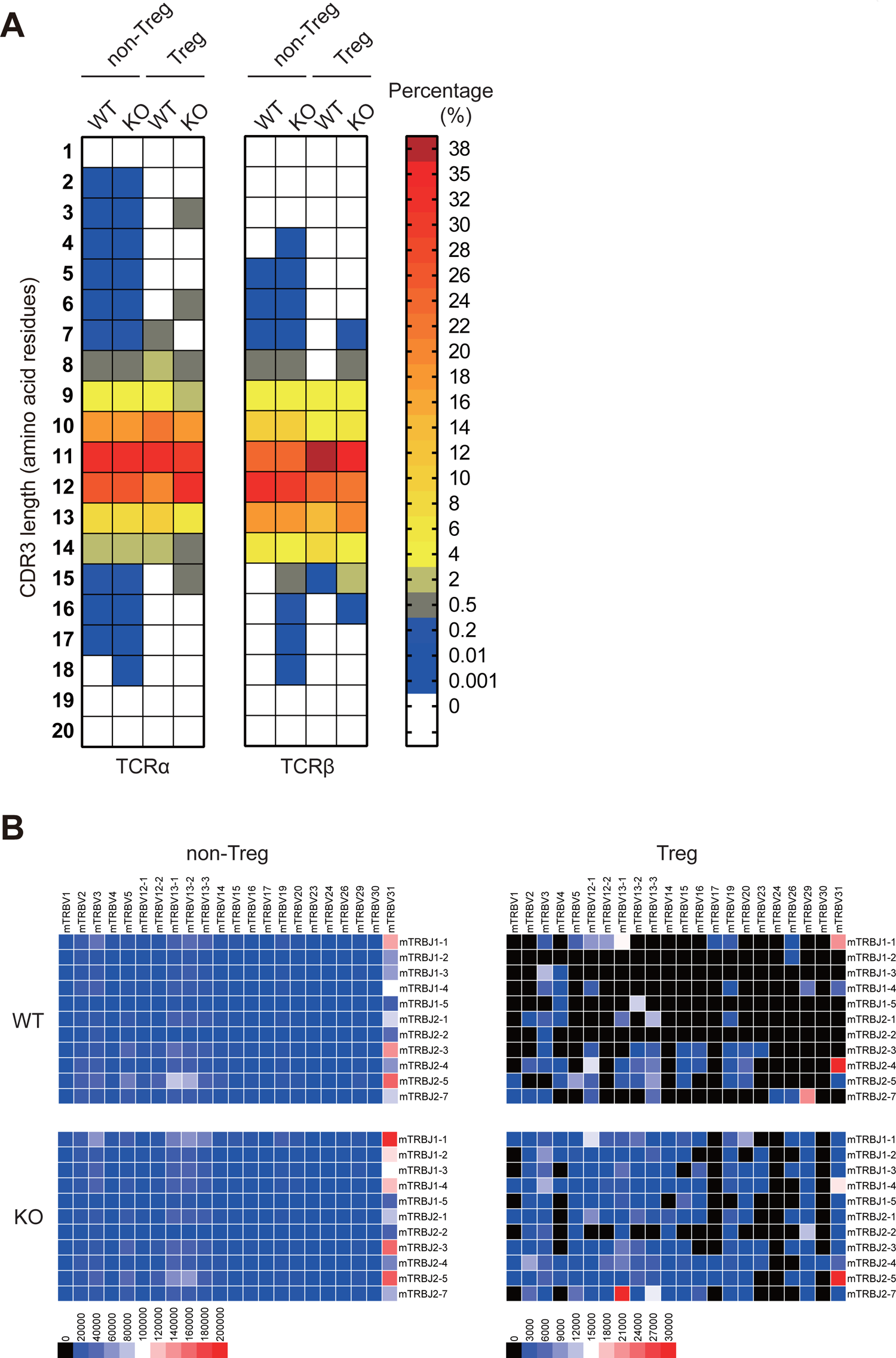
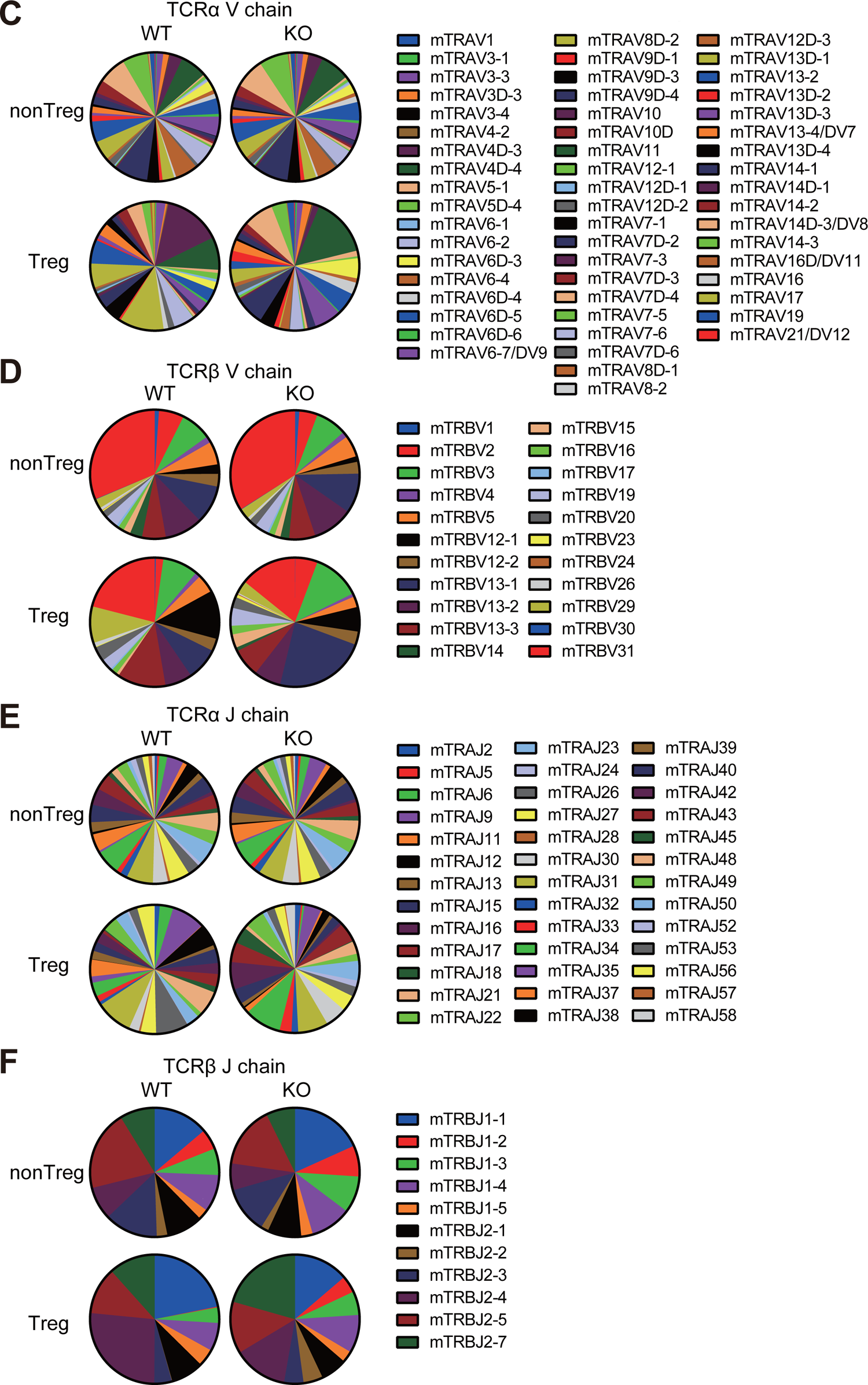
Analysis of TCR repertoires of CIC-deficient CD4^+^ SP thymocytes. **(A-E)** Sequencing analysis of *Tcra* and *Tcrb* mRNAs in thymic non-Treg (CD4^+^CD8^-^GFP^-^) and Treg (CD4^+^CD8^-^CD25^+^GFP^+^) cells from *Foxp3-GFP;Cic^f/f^* (WT) and *Foxp3-GFP;Cic^f/f^;Vav1-Cre* (KO) mice. Heat map showing the distribution of CDR3 amino acid length in the TCRa and TCRb chains **(A)**. Heat map showing the diversity of V and J segment combinations in the TCRβ chain **(B)**. Pie-chart graphs for the usage frequency of V segments of TCRα **(C)** and TCRβ **(D)** chains, and J segments of TCRα **(E)** and TCRβ **(F)** chains.

**Supplemental Figure 4.**
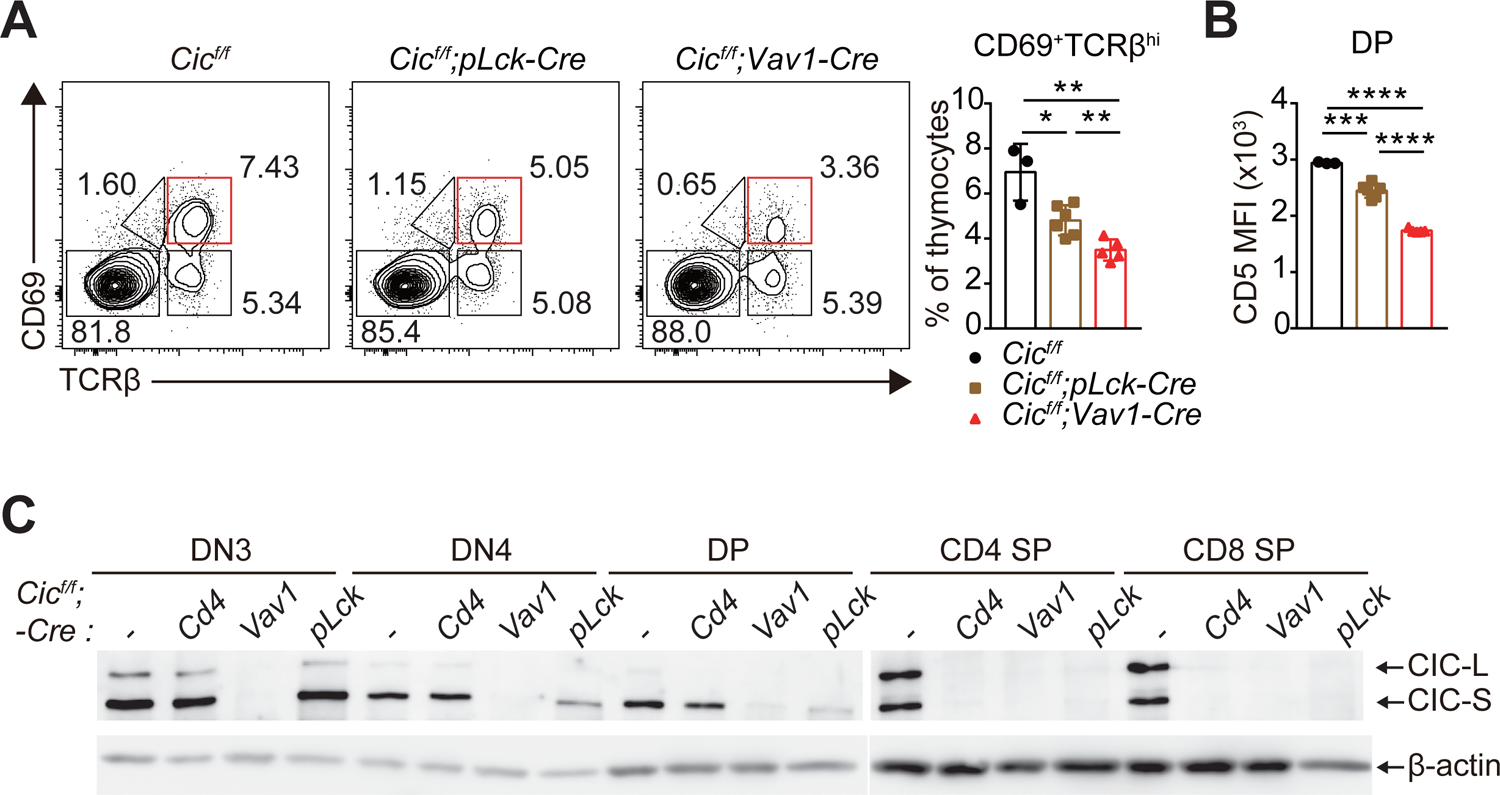
Analysis of thymocytes in *Cic^f/f^;pLck-Cre* mice. **(A and B)** Flow cytometry of thymocytes from 7-week-old *Cic^f/f^* (N = 3), *Cic^f/f^;pLck-Cre* (N = 6), and *Cic^f/f^;Vav1-Cre* (N = 5) mice for the frequency of CD69^+^TCRb^hi^ thymocytes **(A)** and CD5 levels in DP thymocytes **(B)**. The CD69^+^TCRb^hi^ cell population is highlighted with a red box in the FACS plots. Bar graphs show data as means with SEM. *P < 0.05, **P < 0.01, ***P < 0.001, and ****P < 0.0001. **(C)** Western blotting for levels of CIC in DN3, DN4, ISP, DP, CD4^+^ SP, and CD8^+^ SP cells from *Cic^f/f^*, *Cic^f/f^; Cd4-Cre*, *Cic^f/f^; Vav1-Cre*, and *Cic^f/f^;pLck-Cre* mice. Lin^-^ gated DN3, DN4, ISP, DP, CD4^+^ SP, and CD8^+^ SP cells were sorted from mice of each genotype.

**Supplemental Figure 5.**
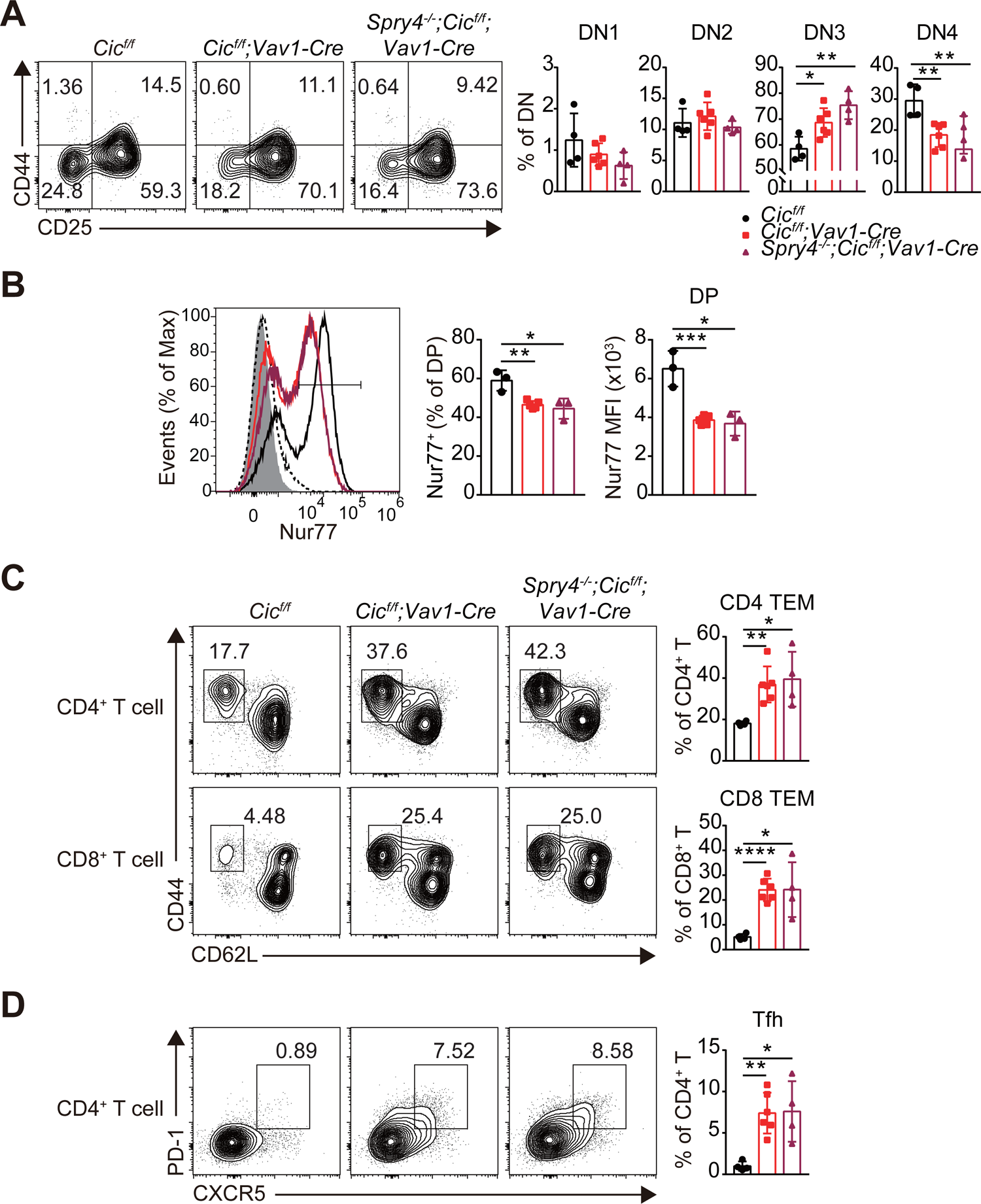
Analysis of T cell subsets in the thymus and spleen of *Spry4^-/-^;Cic^f/f^;Vav1-Cre* mice. **(A)** Flow cytometry of thymocytes from 7-week-old *Cic^f/f^* (N = 4), *Cic^f/f^;Vav1-Cre* (N = 6), and *Spry4^-/-^;Cic^f/f^;Vav1-Cre* (N = 4) mice for the frequency of DN1–4 subsets. **(B)** FACS analysis of Nur77 expression in DP thymocytes from 7-week-old *Cic^f/f^* (N = 3), *Cic^f/f^;Vav1-Cre* (N = 5), and *Spry4^-/-^;Cic^f/f^;Vav1-Cre* (N = 3) mice. Freshly isolated thymocytes were treated with plate-coated anti-CD3 (5 μg/ml) and anti-CD28 (10 μg/ml) for 2 h and then subjected to flow cytometry. Representative histograms of Nur77 expression in DP thymocytes of *Cic^f/f^* (black line), *Cic^f/f^;Vav1-Cre* (red line), and *Spry4^-/-^;Cic^f/f^;Vav1-Cre* (burgundy line) mice overlaid with isotype control (gray shaded) and unstimulated control (dotted line) histograms (left), the frequency of Nur77^+^ DP cells (middle), and Nur77 MFI of DP thymocytes (right) are presented. **(C and D)** Flow cytometry of the frequency of CD44^hi^CD62L^lo^ effector memory T (TEM) **(C)** and Tfh **(D)** cells in the spleen of 7-week-old *Cic^f/f^* (N = 4), *Cic^f/f^;Vav1-Cre* (N = 6), and *Spry4^-/-^;Cic^f/f^;Vav1-Cre* (N = 4) mice. Bar graphs show data as means with SEM. *P < 0.05, **P < 0.01, ***P < 0.001, and ****P < 0.0001.

**Table S1.**
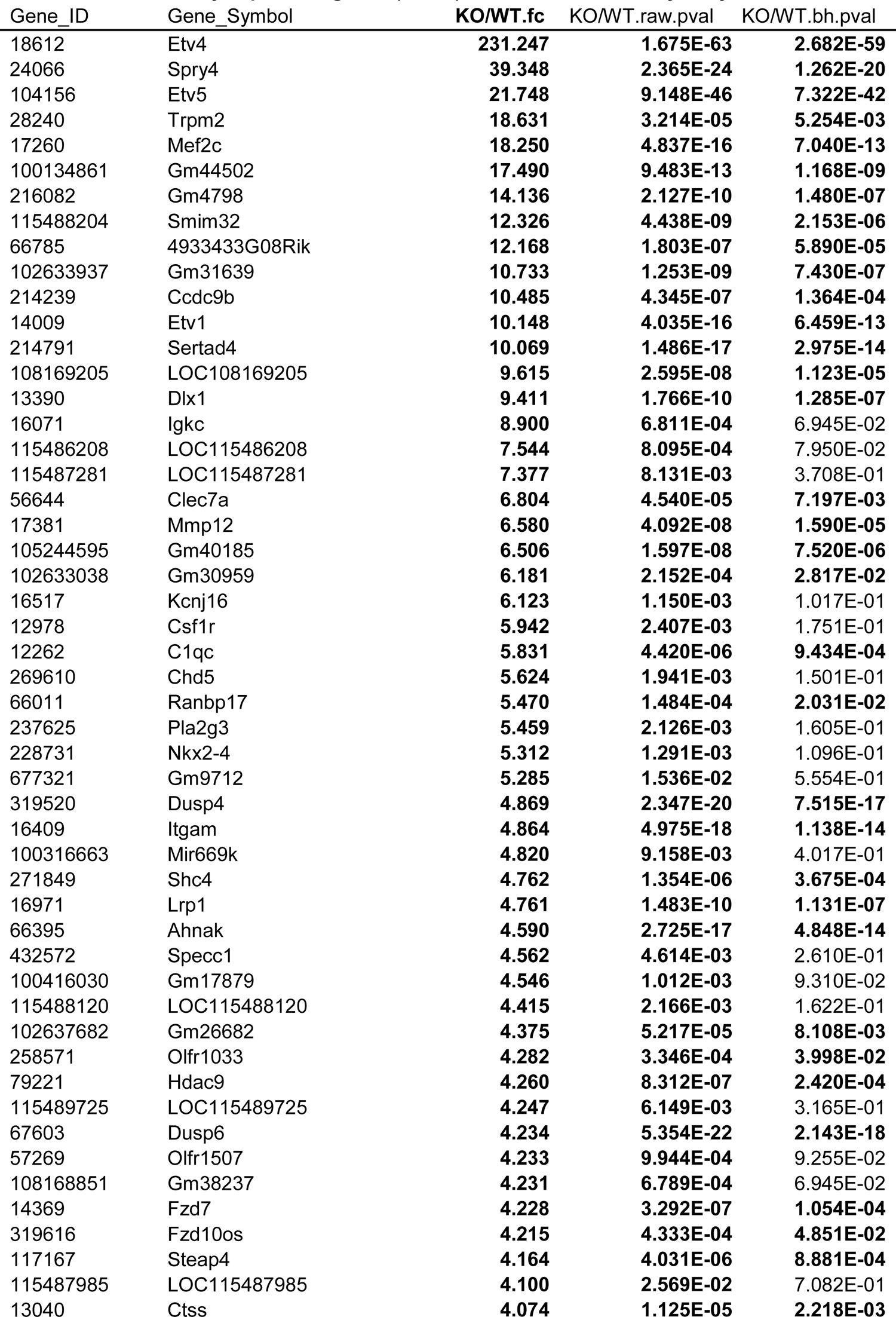
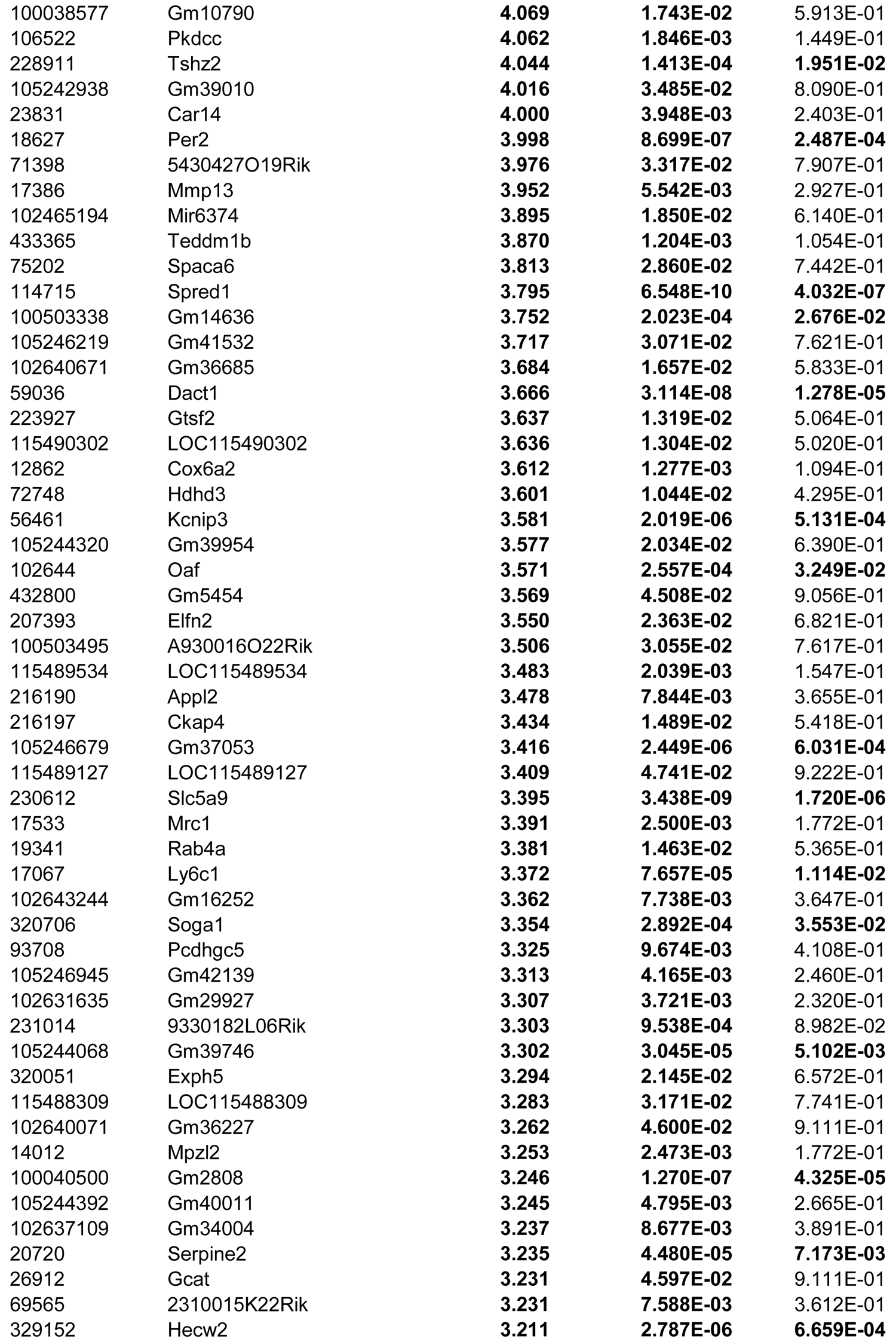
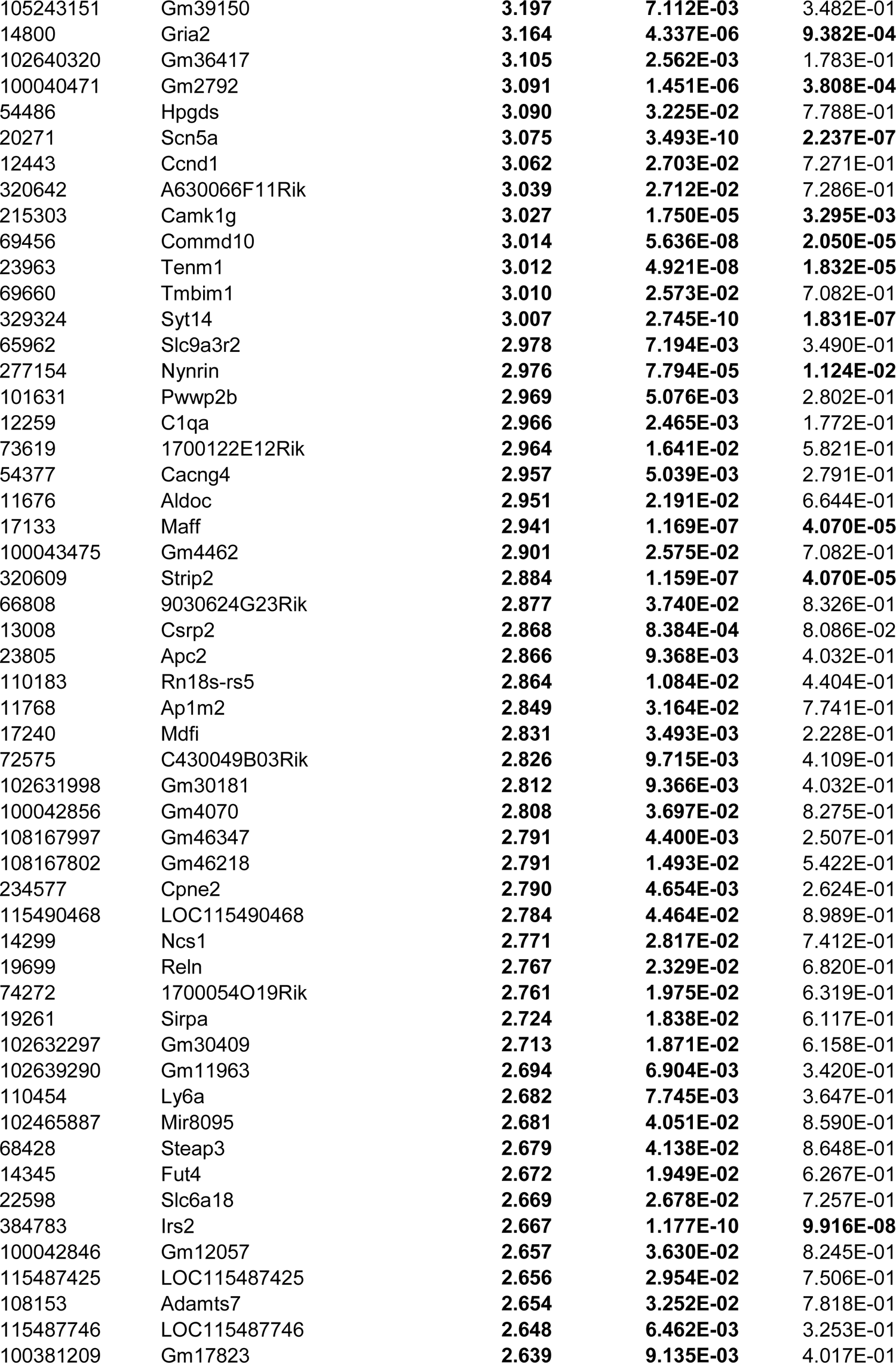
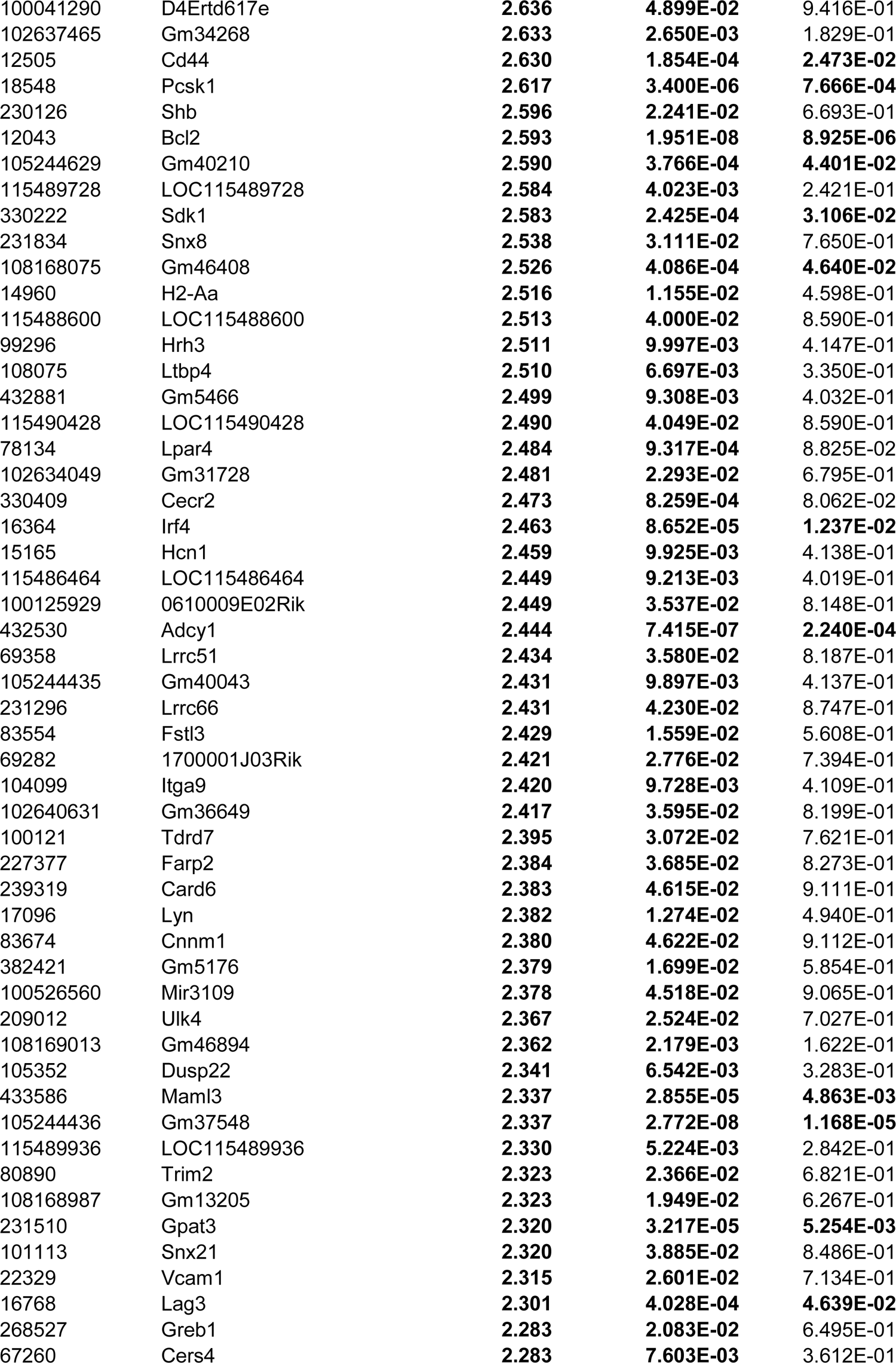
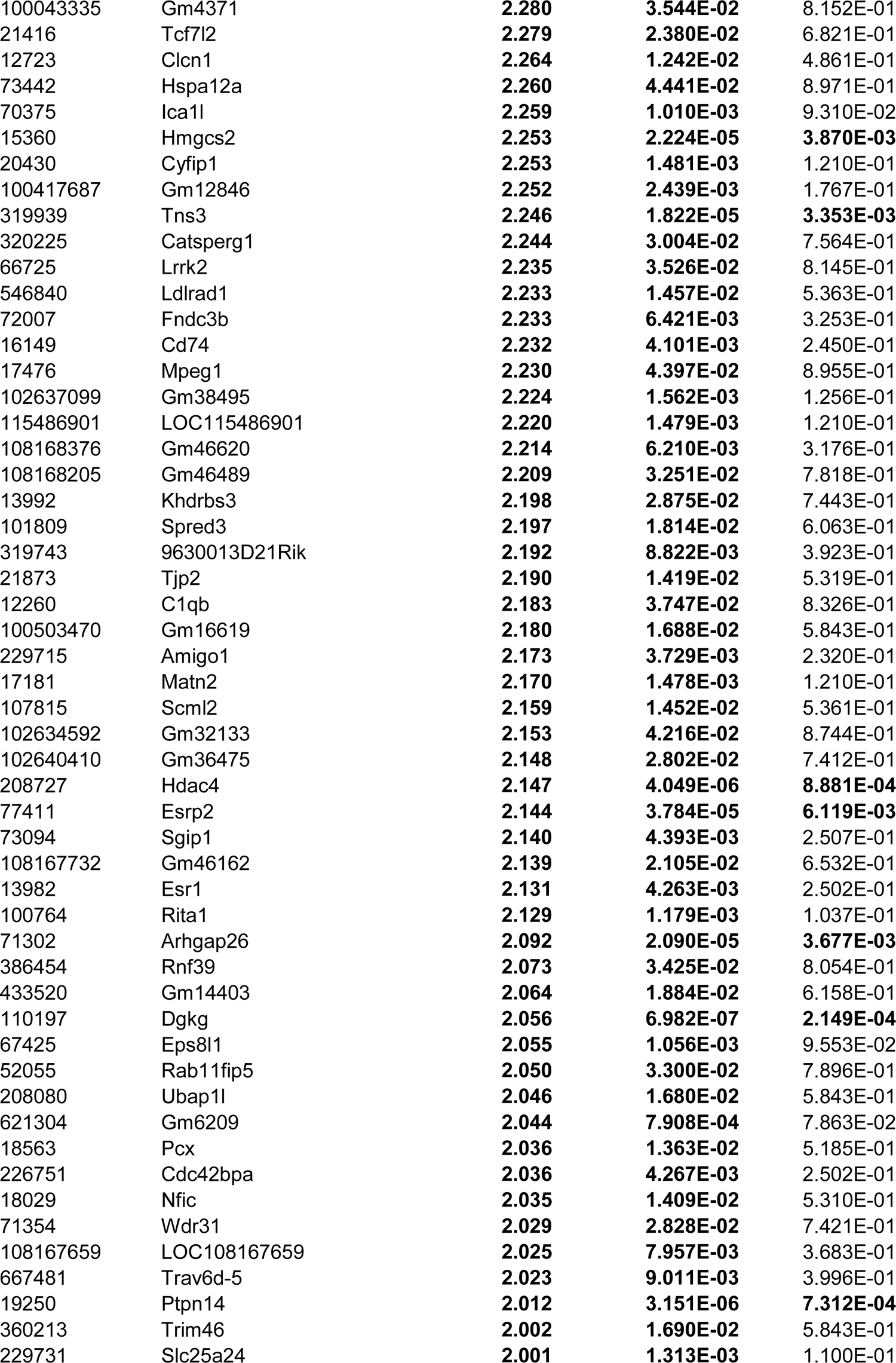
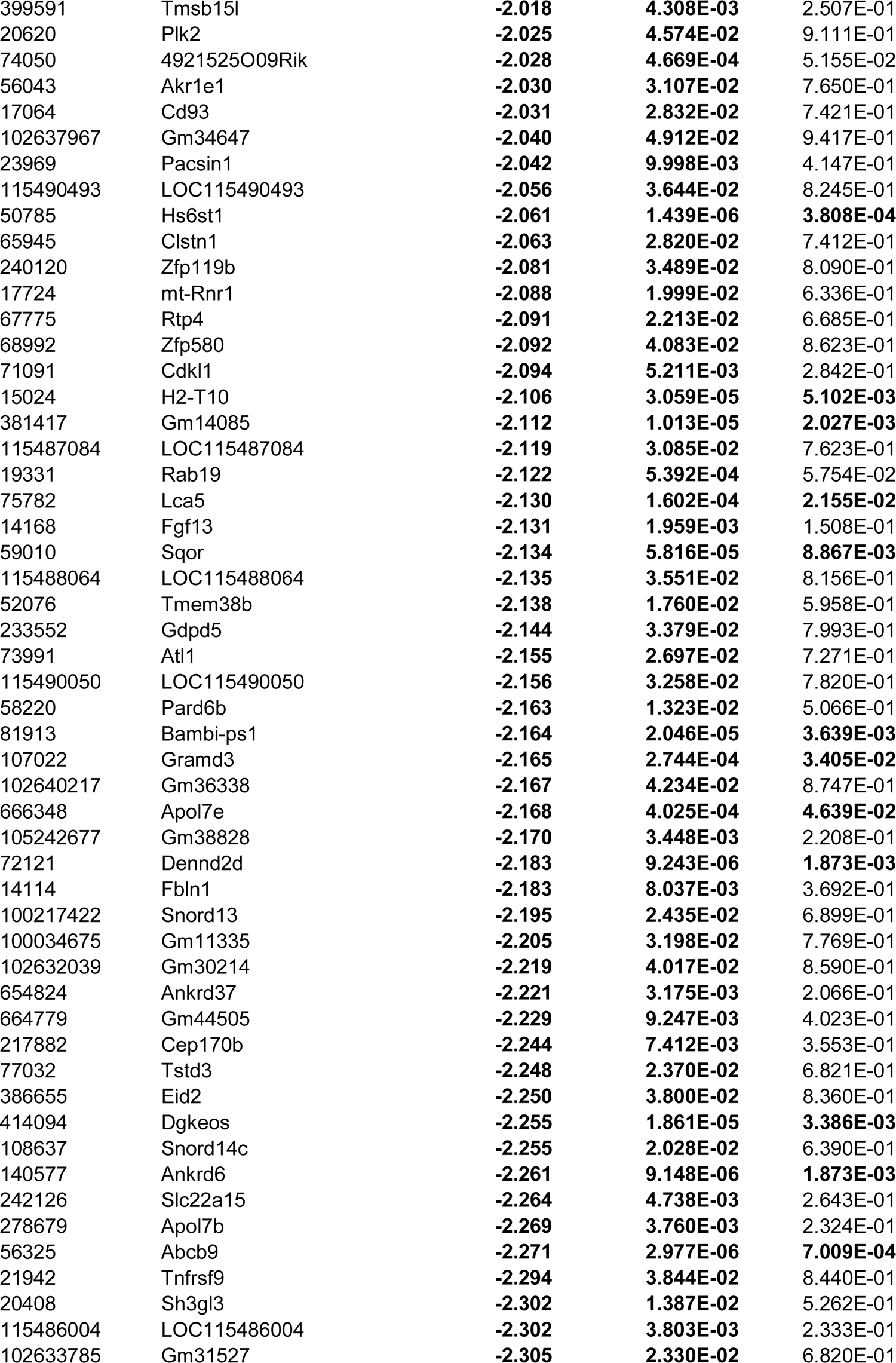
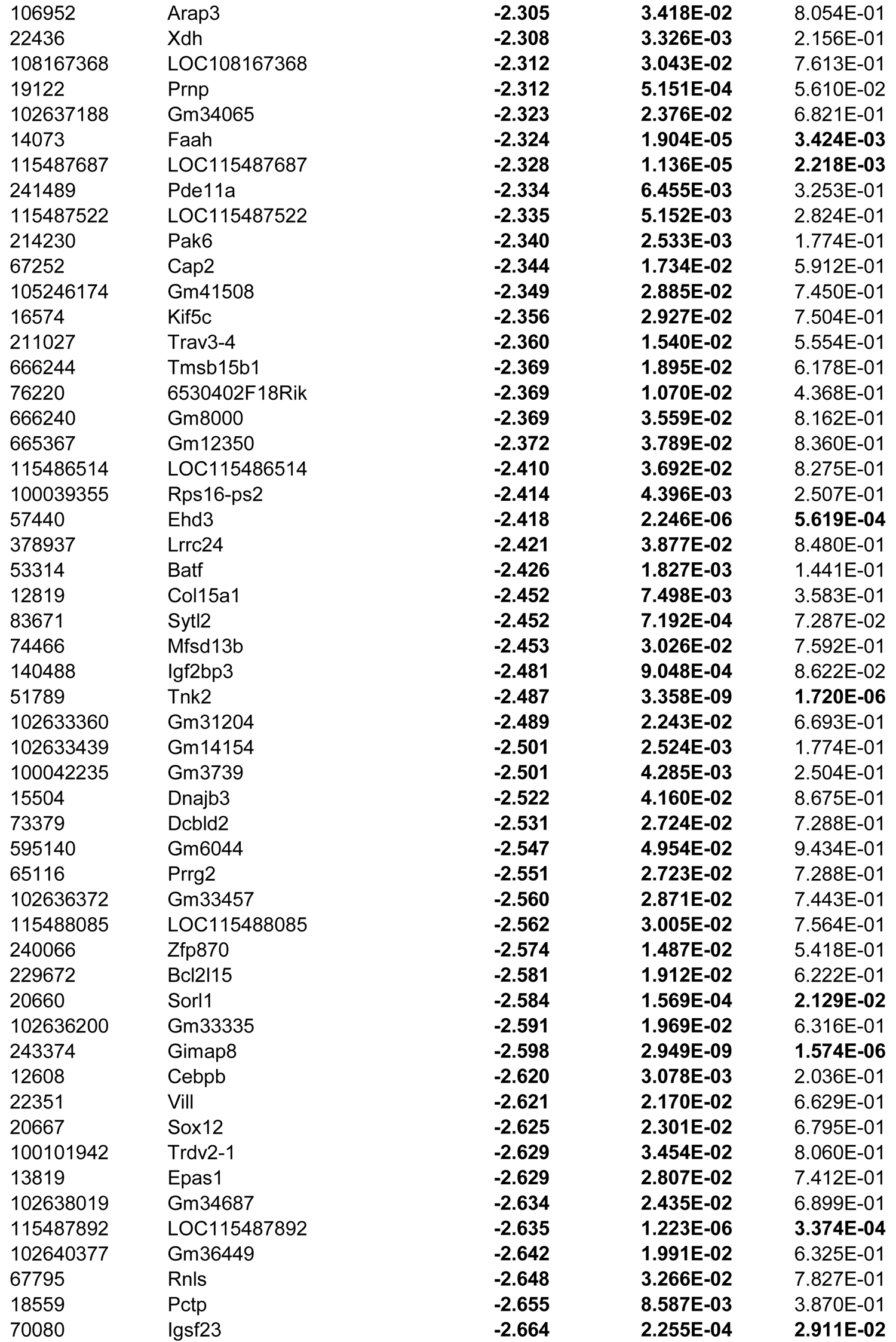
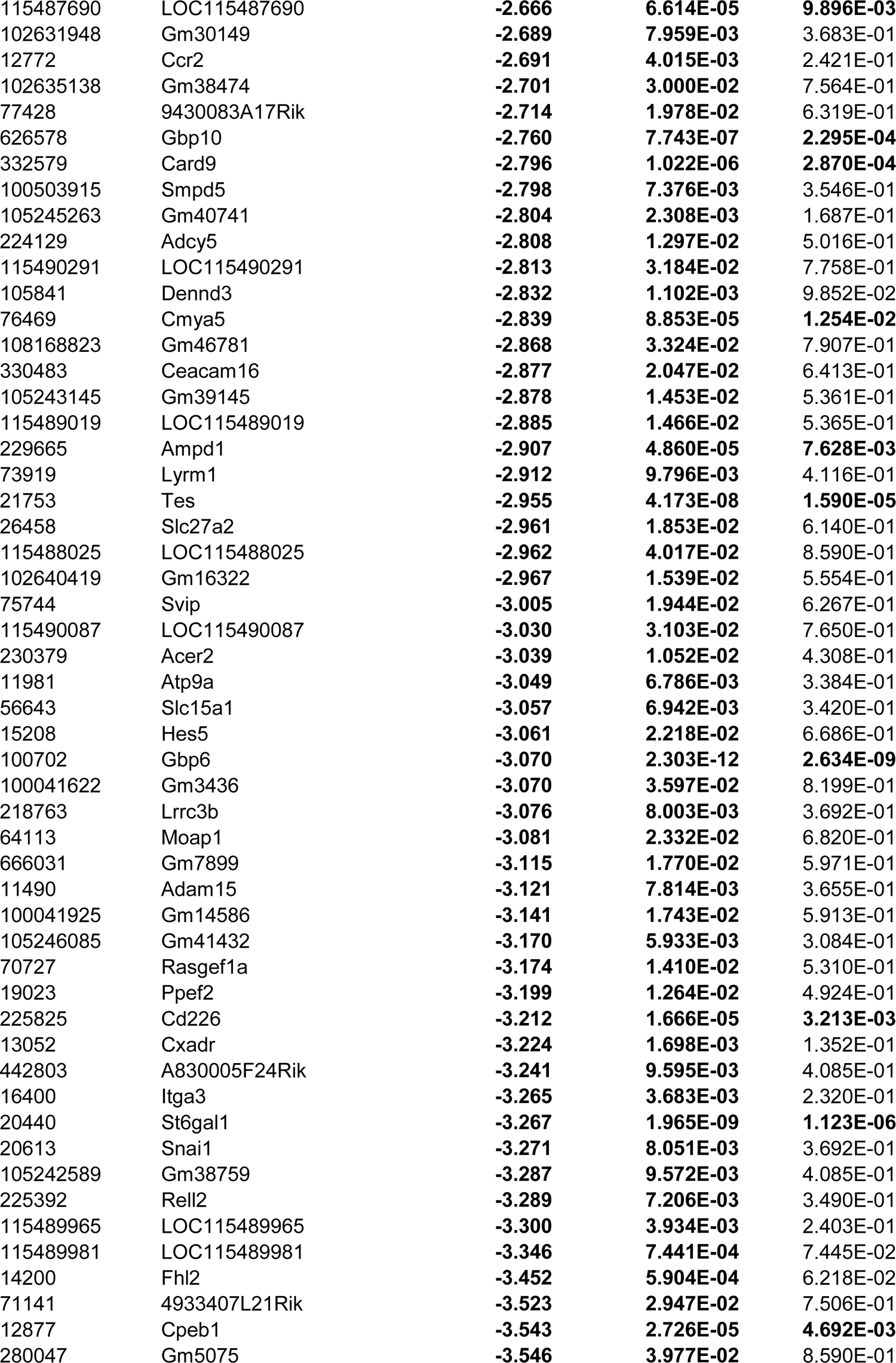
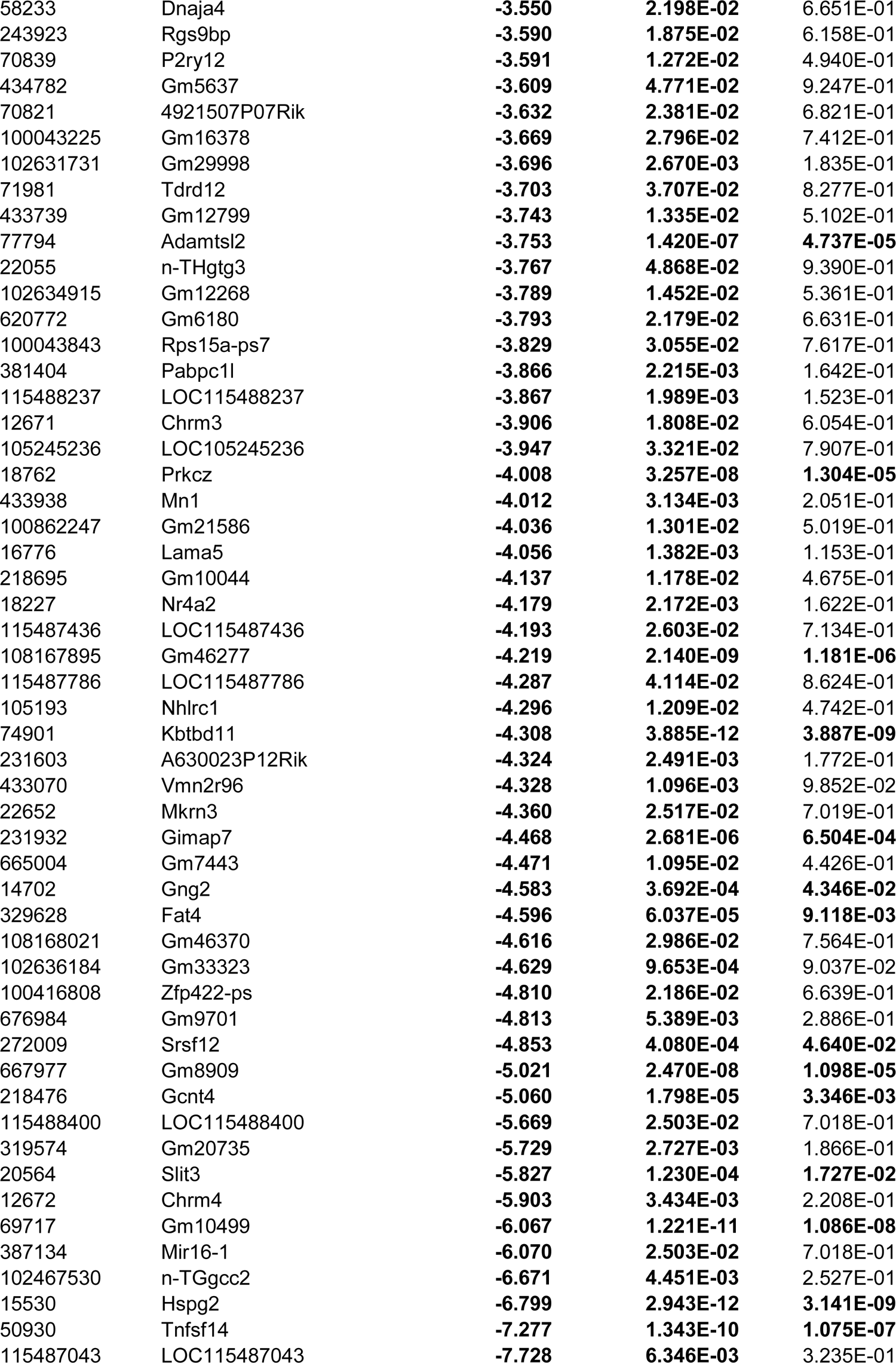
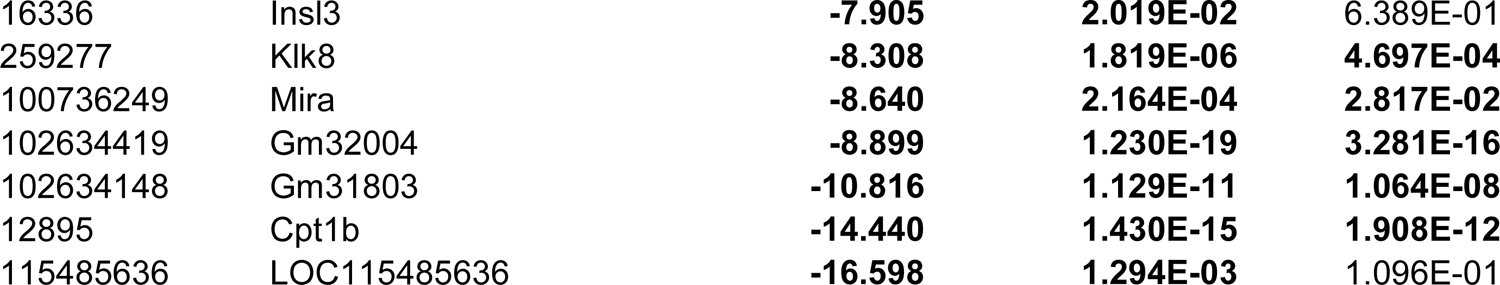
Differentially expressed genes (DEGs) in Cic-deficient DP thymocytes

**Table S2-1.**
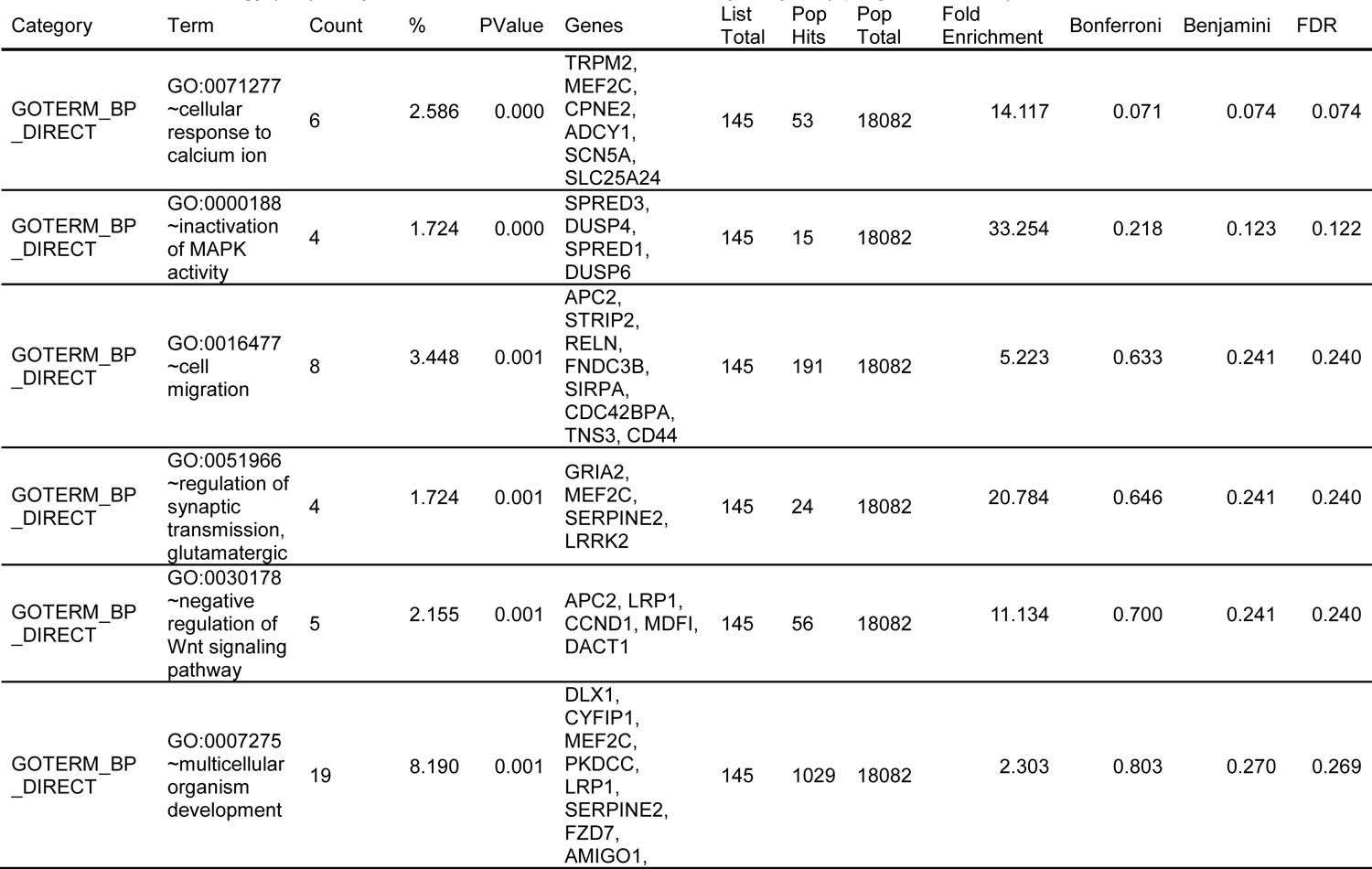
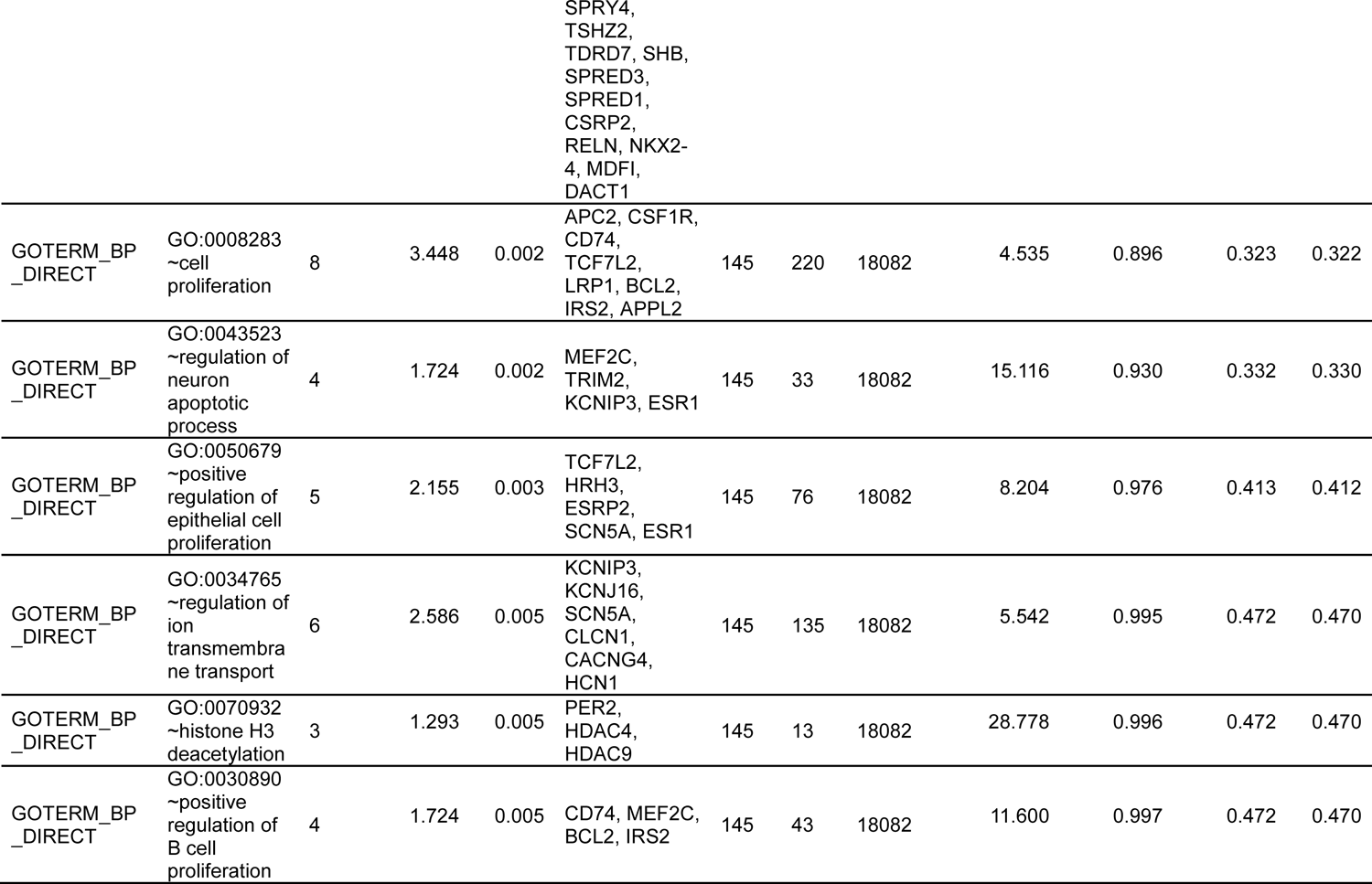
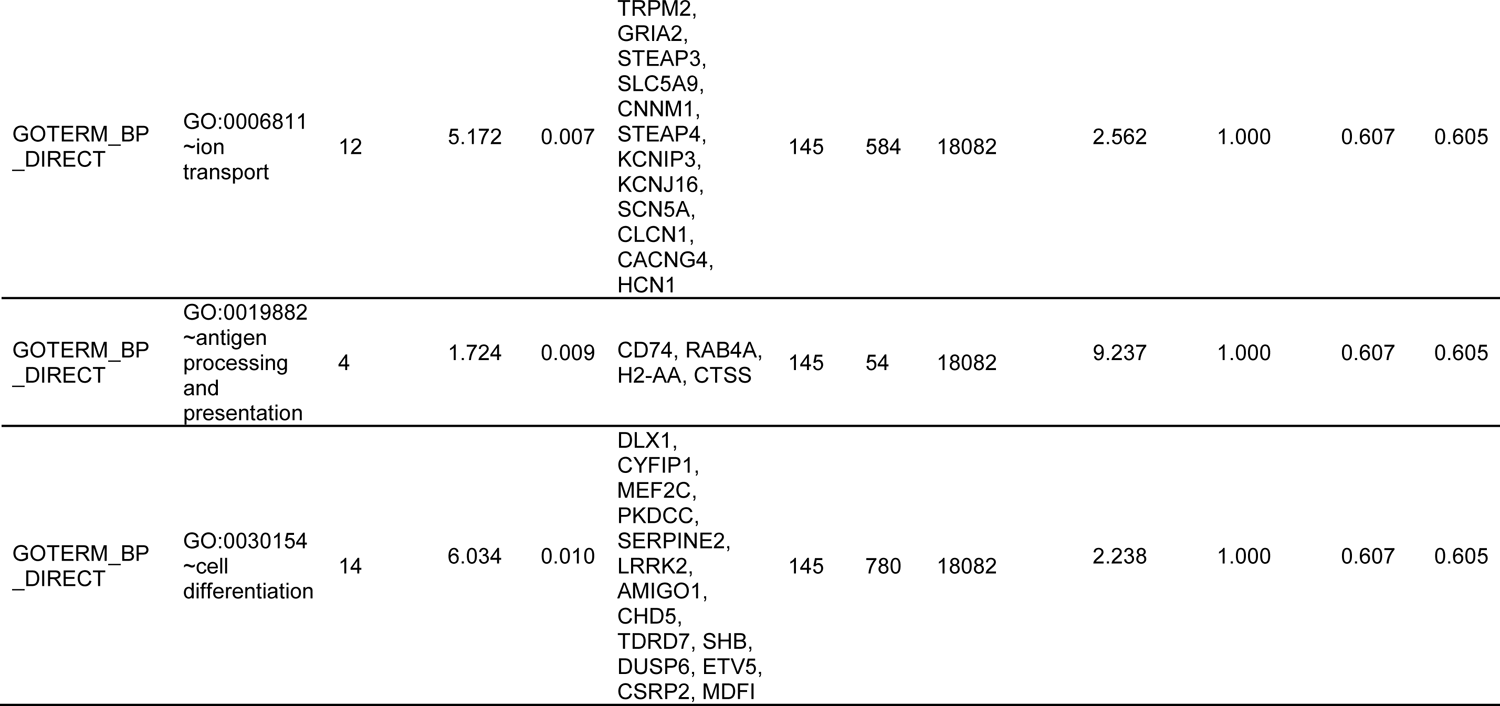
Gene Ontology (GO) analysis of the DEGs in Cic-deficient DP thymocytes (upregulated DEGs)

**Table S2-2.** Gene Ontology (GO) analysis of the DEGs in Cic-deficient DP thymocytes (downregulated DEGs)

| Category | Term | Count | % | PValue | Genes | List Total | Pop Hits | Pop Total | Fold Enrichment | Bonferroni | Benjamini | FDR |
| --- | --- | --- | --- | --- | --- | --- | --- | --- | --- | --- | --- | --- |
| GOTERM_BP_DIRECT | GO:0007613<br>~memory | 6 | 3.141 | 0.000 | CEBPB, ITGA3, PLK2, PAK6, FGF13, KLK8 | 113 | 83 | 18082 | 11.568 | 0.114 | 0.121 | 0.121 |
| GOTERM_BP_DIRECT | GO:0007155<br>~cell adhesion | 10 | 5.236 | 0.003 | LAMA5, COL15A1, CXADR, ADAM15, CD93, ITGA3, CLSTN1, CD226, FAT4, HES5 | 113 | 485 | 18082 | 3.299 | 0.905 | 1.000 | 1.000 |
| GOTERM_BP_DIRECT | GO:0016477<br>~cell migration | 6 | 3.141 | 0.007 | LAMA5, RASGEF1A, TNK2, SNAI1, SORL1, PRKCZ | 113 | 191 | 18082 | 5.027 | 0.993 | 1.000 | 1.000 |
| GOTERM_BP_DIRECT | GO:0042832<br>~defense response to protozoan | 3 | 1.571 | 0.015 | GBP6, GBP10, BATF | 113 | 30 | 18082 | 16.002 | 1.000 | 1.000 | 1.000 |
| GOTERM_BP_DIRECT | GO:0045408<br>~regulation of interleukin-6 biosynthetic process | 2 | 1.047 | 0.018 | CEBPB, CARD9 | 113 | 3 | 18082 | 106.678 | 1.000 | 1.000 | 1.000 |
| GOTERM_BP_DIRECT | GO:0006897<br>~endocytosis | 5 | 2.618 | 0.026 | SH3GL3, TNK2, SORL1, ATP9A, PACSIN1 | 113 | 181 | 18082 | 4.420 | 1.000 | 1.000 | 1.000 |
| GOTERM_BP_DIRECT | GO:0070885<br>~negative regulation of calcineurin-NFAT signaling cascade | 2 | 1.047 | 0.031 | PRNP, CMYA5 | 113 | 5 | 18082 | 64.007 | 1.000 | 1.000 | 1.000 |
| GOTERM_BP_DIRECT | GO:0006869<br>~lipid transport | 4 | 2.094 | 0.032 | APOL7B, PCTP, SORL1, APOL7E | 113 | 111 | 18082 | 5.766 | 1.000 | 1.000 | 1.000 |
| GOTERM_BP_DIRECT | GO:0030198<br>~extracellular matrix organization | 4 | 2.094 | 0.034 | LAMA5, ADAMTSL2, FBLN1, HSPG2 | 113 | 114 | 18082 | 5.615 | 1.000 | 1.000 | 1.000 |
| GOTERM_BP_DIRECT | GO:0007157<br>~heterophilic cell-cell adhesion via plasma membrane cell adhesion molecules | 3 | 1.571 | 0.041 | CXADR, CD226, FAT4 | 113 | 52 | 18082 | 9.232 | 1.000 | 1.000 | 1.000 |
| GOTERM_BP_DIRECT | GO:0007197<br>~adenylate cyclase-inhibiting G-protein coupled acetylcholine receptor signaling pathway | 2 | 1.047 | 0.043 | CHRM3, CHRM4 | 113 | 7 | 18082 | 45.719 | 1.000 | 1.000 | 1.000 |
| GOTERM_BP_DIRECT | GO:0007207<br>~phospholipase C-activating G-protein coupled acetylcholine<br>receptor<br>signaling<br>pathway | 2 | 1.047 | 0.043 | CHRM3, CHRM4 | 113 | 7 | 18082 | 45.719 | 1.000 | 1.000 | 1.000 |
| GOTERM_BP<br>_DIRECT | GO:0001973<br>~adenosine<br>receptor<br>signaling<br>pathway | 2 | 1.047 | 0.049 | P2RY12,<br>ADCY5 | 113 | 8 | 18082 | 40.004 | 1.000 | 1.000 | 1.000 |
| GOTERM_BP<br>_DIRECT | GO:0072006<br>~nephron<br>development | 2 | 1.047 | 0.049 | ITGA3, FAT4 | 113 | 8 | 18082 | 40.004 | 1.000 | 1.000 | 1.000 |
| GOTERM_BP<br>_DIRECT | GO:0015833<br>~peptide<br>transport | 2 | 1.047 | 0.049 | SLC15A1,<br>ABCB9 | 113 | 8 | 18082 | 40.004 | 1.000 | 1.000 | 1.000 |

**Table S3.**
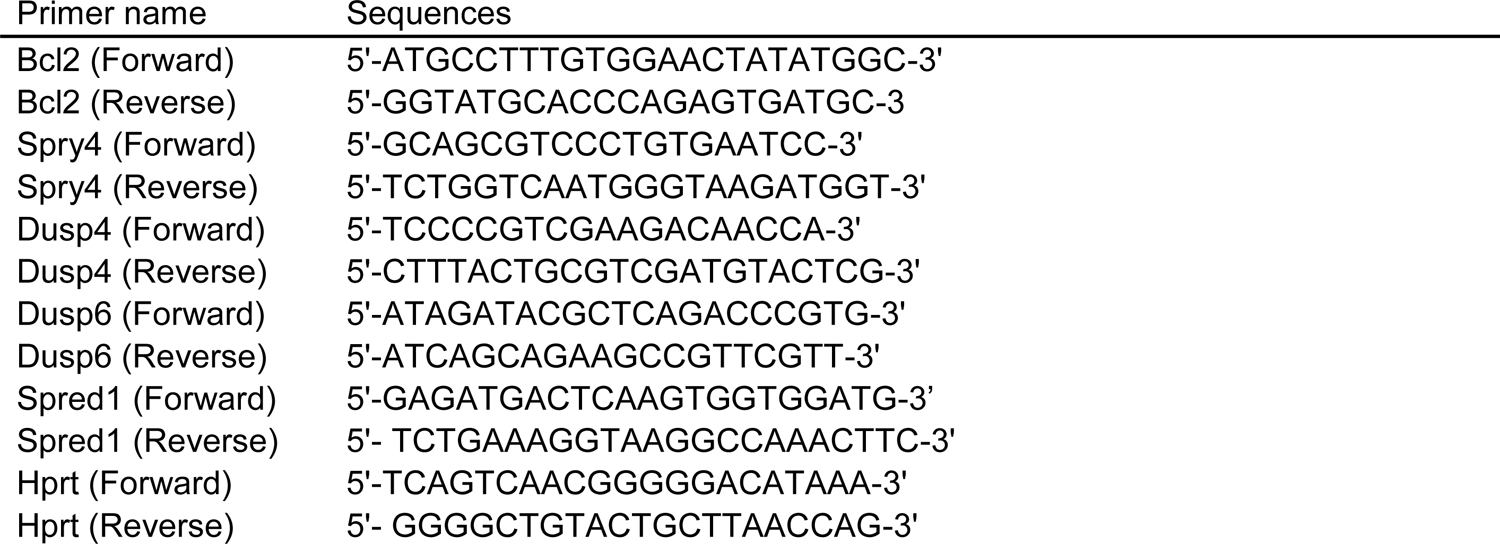
Oligonucleotide sequences used for qRT-PCR

